# Early synaptic changes and reduced brain connectivity in PD-like mice with depressive phenotype

**DOI:** 10.1101/2025.04.24.650442

**Authors:** Lluis Miquel-Rio, Judith Jericó-Escolar, Unai Sarriés-Serrano, Claudia Yanes-Castilla, María Torres-López, Uxia Argibay, Verónica Paz, Carme Casal, Emma Muñoz-Moreno, Xavier López-Gil, Analia Bortolozzi

## Abstract

Anxiety and depression are common in Parkinson’s disease (PD), affecting quality of life. Aggregates of α-synuclein (α-Syn) are found in serotonergic (5-HT) raphe nuclei early in the disease, but their relationship to brain changes is unclear. We investigated synaptic plasticity, neuronal activity, and functional magnetic resonance imaging (fMRI)-based brain connectivity in a PD-like mouse model with depressive phenotype. AAV-induced human α-Syn accumulation in raphe 5-HT neurons causes progressive synaptic pathology in interconnected brain regions. This is marked by lower MAP-2, PSD95 and higher SV2A, VAMP2, which are key to synaptic structure and function, as confirmed in human brain tissue samples. Abnormalities in Egr-1-dependent neuronal activity and region-specific differences in resting-state functional brain activity were also detected eight weeks post-AAV infusion, before neurodegeneration. This provides evidence for synaptic and fMRI markers associated with α-Syn pathology in emotional brain circuits, and has translational importance for identifying PD patients at risk for depression.

## INTRODUCTION

Parkinson’s disease (PD) is the most prevalent neurodegenerative motor disorder worldwide, with resting tremor, rigidity, bradykinesia and postural instability being the main symptoms of the disease ^1,2^. However, growing evidence suggests that PD is a disorder with numerous non-motor symptoms that affect multiple, non-dopaminergic neuronal populations aside from the dopaminergic (DA) system ^3,4^. Among mental health disorders, depression is one of the major non-motor symptoms of PD, with approximately 40–50% of patients with PD suffering from depressive disorder ^5–7^. Although depression may be secondary to progressive and disabling motor symptoms, a significant number of PD patients present comorbid depression early in the prodromal phase, and many have somewhat more rapid disease progression ^8–10^. This suggests that depression may influence primary motor impairments or signal more severe forms of PD. Therefore, it is not surprising that converging evidence suggests that a first occurrence of depression significantly increases the risk of developing PD later in life.

Considerable efforts have been devoted to elucidate the neural basis of depression in PD (D-PD). Postmortem studies suggest that pathological processes may occur early in cortical and limbic regions linked to emotional control ^11–13^, providing insight into the pathological mechanisms of depression in the pre-motor PD phase. Indeed, neurotransmitter imaging studies have shown abnormal serotonin (5-HT) innervation and neurotransmission in different brain regions including the dorsolateral prefrontal cortex (dlPFC), orbitofrontal cortex, anterior cingulate cortex (ACC), caudate nucleus (CAU), thalamus, and hippocampus (HPC) in D-PD patients ^14–17^. However, other neuroimaging studies also suggest a loss of DA and noradrenaline fibers in the limbic regions in patients with D-PD ^13,18,19^. Abnormal functional activity in the PFC, e.g. dlPFC and ACC, has been characterized as a critical feature in the pathophysiology of depressive disorders, as well as in the favorable outcomes of new antidepressant strategies ^20–23^. Of note, the PFC is strongly innervated by raphe 5-HT neurons ^24,25^ and 5-HT dysfunction is linked to depressive disorder, also in PD ^16,26^. Consistent with these data, several structural and functional neuroimaging studies have shown pronounced changes in the PFC networks in D-PD patients that differ from those of patients with PD alone ^27–29^. Previous studies using 2-[^18^F]-fluoro-2-deoxy-D-glucose and positron emission tomography (PET) showed hypo-metabolism in the orbitofrontal cortex and ACC ^30,31^, suggesting that D-PD may be associated with orbital-inferior frontal lobe dysfunction. In addition, using magnetic resonance imaging (MRI), cortical thinning and white matter abnormalities have been reported in prefrontal, temporal and some limbic regions in patients with D-PD ^32–34^. Reduced bilateral HPC and amygdala (Amg) volumes are other neuroanatomical correlates of depressive symptoms in PD ^35,36^.

More recently, it has been proposed that abnormal cortical-limbic circuit activity and connectivity, which suggest altered higher-order control of negative mood, is a possible neural mechanism of D-PD. Resting-state functional MRI (rs-fMRI) analysis showed reduced functional connectivity between the ACC and the right tempo-parietal junction, as well as between the posterior cingulate cortex and the insula, and between the superior parietal lobule and the PFC in patients with D-PD, distinguishing them from those PD patients without depression ^28,37–39^. Other connectome studies have also reported a decrease in functional connectivity within the cortical-limbic networks, together with an increase in connectivity between limbic regions, e.g. increased functional connectivity between amygdala with the bilateral mediodorsal thalamus, in patients with D-PD ^37,40,41^. These data suggest that the salient circuitry involved in emotion recognition, and negative emotions in particular, may be impaired in comorbid depressive symptoms in PD, and the underlying pathological mechanisms await further elucidation.

The causes of PD are unclear, but the build-up of the α-synuclein protein (α-Syn) is a key part of the disease process ^16,42^. α-Syn is expressed in all neuronal types, where it modulates axonal transport, vesicle dynamics and, ultimately synaptic plasticity ^43,44^. Its overexpression in mouse raphe 5-HT neurons causes an increase in phosphorylation and aggregation of α-Syn protein, together with progressive accumulation of α-Syn in 5-HT-positive fibers in cortico-limbic regions, deficits in the brain-derived neurotrophic factor (BDNF) expression and reduced 5-HT neurotransmission, resulting in a depressive-like phenotype ^45^. Of note, depression has been associated with elevated plasma α-Syn levels ^46^, and we found higher levels of phospho-α-Syn in postmortem dlPFC and CAU samples from patients with depression, comparable to those of early-stage PD patients (Braak 2-3) ^47^, which seems significant given the prevalence of depression in PD. Here, we combined rs-fMRI with synaptic and cellular activity mapping in the D-PD-like mouse model ^45^ to probe how α-Syn accumulation in the brain 5-HT system leads to synaptic and connectivity dysfunctions in brain areas controlling mood and emotional processes. We found altered synaptic marker density, increased cellular activity, and reduced rs-fMRI coupling of medial PFC (mPFC), caudate-putamen (CPu), and HPC with their cortico-subcortical targets dependent on changes in 5-HT neuroplasticity associated with α-Syn pathology. Changes in synaptic markers were confirmed in post-mortem samples from PD patients. These findings may be useful in facilitating a better understanding of the potential mechanism underlying depression in PD.

## RESULTS

### D-PD-like mouse model with h-α-Syn overexpression in raphe 5-HT neurons

First, we confirmed that expression of the wild-type human α-Syn transgene (h-α-Syn) in 5-HT raphe neurons results in a progressive increase in h-α-Syn protein levels in the mouse 5-HT ascending system, as previously described ^45^. Here we use a novel AAV2/5-CBh-WPRE3A vector encoding h-α-Syn (denoted AAV-α-Syn, provided by the MJJ Foundation) - previously we used AAV5-CBh-α-Syn ^45^ - infused into the mouse dorsal raphe nucleus (DR). To investigate the time-course of h-α-Syn expression, male and female mice were euthanized at 4 and 8 weeks post-injection. Increased h-α-Syn protein density levels were detected in the rostral-caudal axis of the DR (anteroposterior – AP coordinates from -4.24 to -4.72 mm) of both males and females compared to the respective control group injected with the empty AAV2/5-CBh-WPRE3 vector containing non-coding (null) stuffer DNA (AAV-EV) (p < 0.0001) (**Fig. 1A, 1B, Suppl 1A, 1B**). However, while the maximum increase in h-α-Syn density levels was found in male mice 8 weeks later (∼341% vs. ∼477% compared to control group at 4 and 8 weeks, respectively), no significant differences were found in female mice at 4 and 8 weeks post-AAV-α-Syn injection (∼295% and ∼286%, respectively) (**Fig. 1B, Suppl 1B)**. Further histological analysis was performed to assess whether h-α-Syn protein co-localized with cells positive for the neuronal 5-HT-specific marker tryptophan hydroxylase (TPH). We found progressively significant increases in the number of TPH-positive cells co-localizing with h-α-Syn in the DR of male mice. At 8 weeks post-infusion 82.5±1.3% of TPH-positive cells were positive for h-α-Syn immunoreactivity, compared to 66.4±0.9% detected at 4 weeks post-infusion (p < 0.0001) (**Fig. 1B)**. No significant differences were observed in female mice and we found a similar number of double positive cells for TPH and h-α-Syn in the DR (83.1±1.9% and 84.2±0.9% of double TPH/h-α-Syn-positive cells at 4 and 8 weeks, respectively) (**Suppl 1B).** No loss of TPH-positive cells was observed in either sex, at least in this timeline. As we have found sex-related differences in raphe h-α-Syn accumulation and in behavioral profiles, e.g., stress-related behavior in males vs. anxiety-related behavior in females ^48^, here we focused the next experiments on male mice.

**Figure 1.**
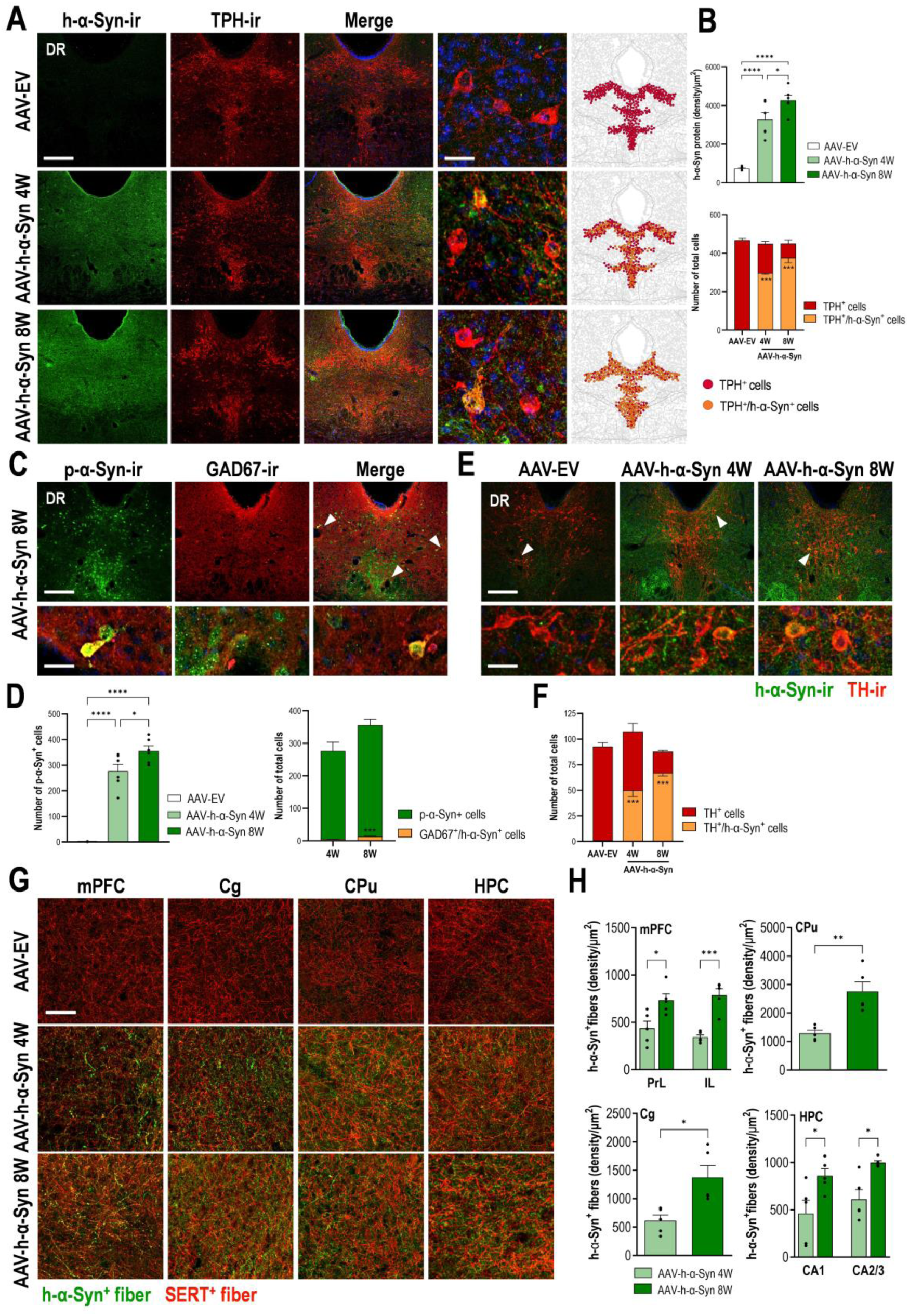
Overexpressing of h-α-Syn in raphe 5-HT neurons in male mice. Male mice received 1 μl of AAV2/5-CBh-WPRE3 construct to drive expression of h-α-Syn (AAV-h-α-Syn) or empty AAV2/5-CBh-WPRE3 vector containing non-coding (null) stuffer DNA (AAV-EV) into dorsal raphe nucleus (DR) and were euthanized at 4 and 8 weeks (W) post-injection. **A**) Representative confocal image showing co-localization of h-α-Syn protein and TPH^+^ cells in the DR of mice after AAV2/5 injection. Scale bar: low: 200 μm, high: 50 μm. On the right, schematic representation of h-α-Syn^+^/TPH^+^ cell density. **B)** Bar chart (top) showing progressive h-α-Syn accumulation in mouse DR. Bar chart (down) showing the proportion of the total number of TPH^+^ cells co-localized with h-α-Syn^+^ (n = 6 mice/group; *p< 0.05, ****p< 0.001 versus AAV-EV or AAV-h-α-Syn 4W mice). **C)** Representative confocal image showing co-localization of p-α-Syn protein and GAD67^+^ cells in the DR of mice at 8W after AAV2/5 injection. Scale bar: low: 200 μm, high: 50 μm. **D)** Bar chart (left) showing progressive accumulation of p-α-Syn protein in mouse DR. Bar chart (right) showing the number of double cells co-localizing GABA67^+^/p-α-Syn^+^ in mouse DR (n = 6 mice/group; *p< 0.05, ****p< 0.001 versus AAV-EV or AAV-h-α-Syn 4W mice). **E)** Representative confocal image showing co-localization of h-α-Syn protein and TH^+^ cells in the DR of mice at 4W and 8W after AAV2/5 injection. Scale bar: low: 200 μm, high: 50 μm. **F)** Bar chart showing the proportion of the total number of TH^+^ cells co-localized with h-α-Syn^+^ (n = 5 mice/group; ***p< 0.001 versus AAV-EV mice). **G)** Representative confocal image showing co-localization of h-α-Syn^+^/SERT^+^ fibers in 5-HT projection brain regions, such as medial prefrontal cortex (mPFC), cingulate cortex (Cg), caudate putamen (CPu), and hippocampus (HPC) of mice at different time points after AAV2/5 injection. Scale bar: 25 μm. **H)** Bar chart showing progressive accumulation of h-α-Syn protein in projection brain areas: prelimbic cortex (PrL), infralimbic cortex (IL), Cg, CPu, and different HPC subfields (n = 5 mice/group; *p< 0.05, **p< 0.01, ***p < 0.001 versus AAV-h-α-Syn 4W mice). Values are presented as mean ± SEM. See Supplementary Fig. 1-3.

In the DR, the five major neuronal classes (in decreasing order of abundance) are 5-HT, DA, GABA, glutamate, and peptidergic neurons ^49^. Therefore, we examined whether the two most important non-5-HT neuronal populations in the DR, GABA and DA neurons, are positive for h-α-Syn or its phosphorylated form at amino-acid serine-129 (p-α-Syn). We found that of the total number of p-α-Syn-positive cells in the DR, a small number were also positive for the GABA GAD67 marker (1.9±0.2% and 3.7±0.3% at 4 and 8 weeks, respectively) (**Fig. 1C, 1D**). In addition, we detected 62.4±1.3% and 76.2±0.9% of double-positive cells for the DA tyrosine hydroxylase (TH) neuron marker and h-α-Syn in more rostral DR regions (AP coordinates -4.24 to -4.36 mm) at 4 and 8 weeks post-injection, respectively, with no change in the total number of TH-positive neurons compared to AAV-EV control group (**Fig. 1E, 1F**).

Next, we analyzed the accumulation of h-α-Syn in efferent brain areas including the mPFC, cingulate cortex (Cg), CPu, and HPC of AAV-EV and AAV-α-Syn injected mice. As previously reported ^45^, we confirmed an abundant and progressive presence of h-α-Syn-positive fibers in the different brain areas analyzed, reaching maximum levels 8 weeks later compared to 4 weeks (p < 0.001) (**Fig. 1G, 1H**). The h-α-Syn signal was exclusively axonal and no h-α-Syn cell bodies were detected, as previously (Alarcón-Arís et al., 2020). In parallel, a significant loss of serotonin transporter-(SERT)-positive axons was observed in all brain areas examined 8 weeks later (p < 0.01; **Suppl 2**). In addition, we analyzed SERT/h-α-Syn-positive fiber density and found values (in descending order of density) of 39.2±2.5%, 30.9±2.7%, 20.1±4.5%, and 15.8±1.5% in CPu > Cg > mPFC > HPC at 8 weeks post-injection, indicating that h-α-Syn protein was transported anterogradely along 5-HT axons towards synaptic terminals. More interestingly, we also detected some TH-positive fibers co-localizing with h-α-Syn at 4 and 8 weeks post-injection. These were examined only in CPu and Cg of AAV-h-α-Syn-injected mice in DR compared to the control group (**Suppl 3A, 3B**). Analysis showed a progressive increase in the number of TH/h-α-Syn-positive fibers (CPu: 5.41±0.9%, 10.5±1.9%; Cg: 12.0 2.7%, 21.3 0.8 at 4 and 8 weeks later, respectively), although to a lesser extent compared to the density of SERT/h-α-Syn-positive fibers in the same brain areas. No significant difference in the TH-positive axon density was observed in the brain areas analysed (**Suppl 3C, 3D).**

### Synaptic pathology in efferent brain regions in the D-PD-like mouse model

In physiological conditions, α-Syn is abundant in the pre-synaptic compartment ^51^. Neuropathological and experimental studies provided evidence that α-Syn aggregation might start at pre-synaptic terminals, inducing synaptic degeneration and subsequently neuronal loss ^43,52^. Since an accumulation of the h-α-Syn protein was detected in efferent brain regions after infusion of AAV-h-α-Syn in DR, we mapped synaptic changes using different pre- and post-synaptic markers in interconnected brain areas. Immunoreactivity for synaptic vesicle glycoprotein 2A (SV2A), an integral glycoprotein found in the membranes of synaptic vesicles in all synaptic terminals and widely distributed throughout the brain, was increased by ∼87.8% in Cg (p < 0.0001) and by ∼82.6% in CPu (p < 0.0001) of AAV-h-α-Syn injected mice compared to control group at 8 weeks’ post-infusion. No changes were observed in other brain areas analyzed (**Fig. 2A, 2B)**. In parallel, synaptophysin (SYP) density, another abundant synaptic vesicle membrane protein comprising ∼10% of the total synaptic protein ^53^, was also increased in Cg (∼25%, p < 0.05) and CPu (∼40%, p < 0.05) of the same AAV-h-α-Syn injected mice compared to control group at 8 weeks post-infusion. Within the groups, a similar SYP density was detected in all other brain regions (**Fig. 2C, 2D**).

**Figure 2.**
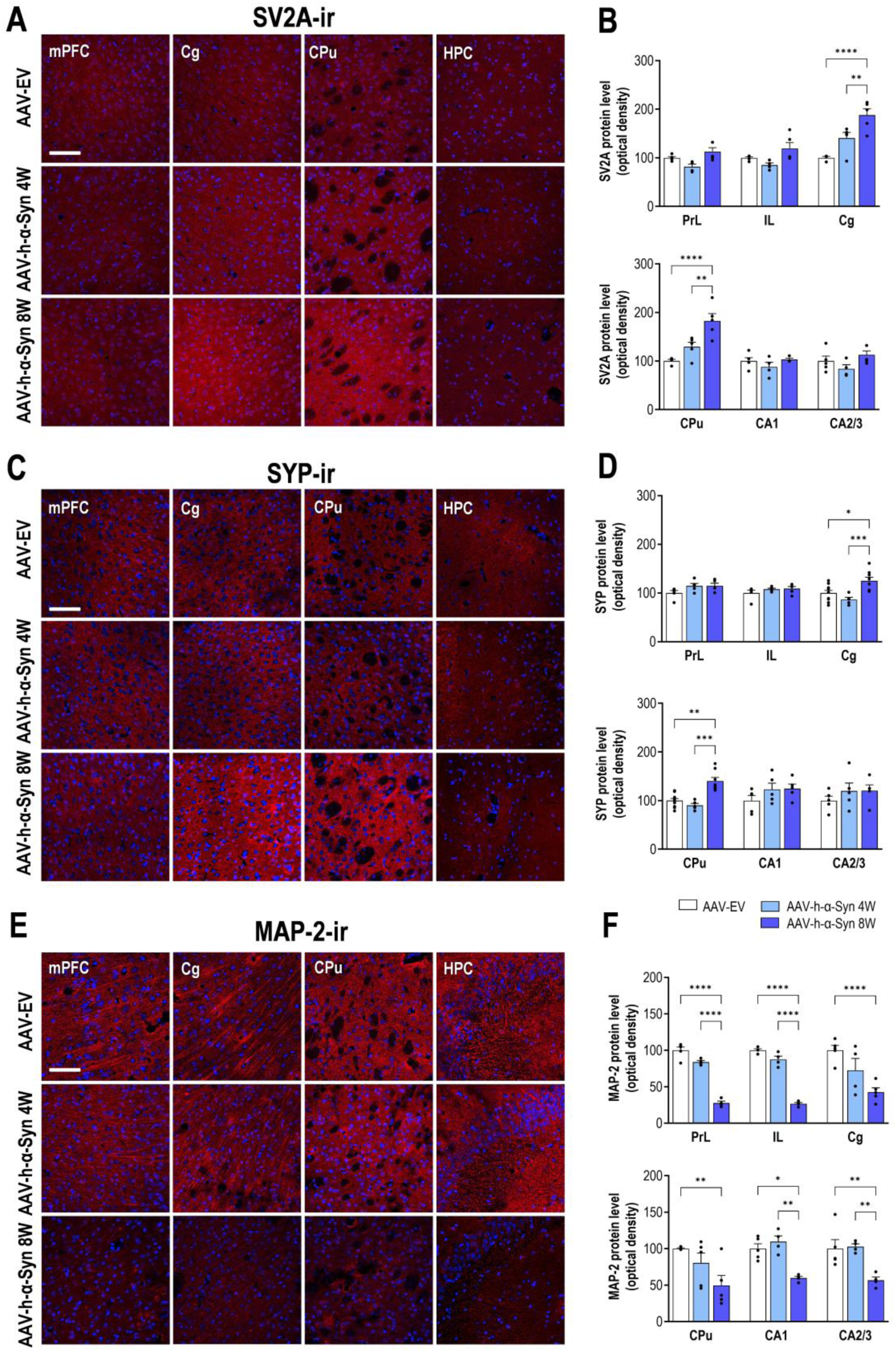
Synaptic protein density in cortical and limbic areas in the D-PD mouse model. Male mice received 1 μl of AAV2/5-CBh-WPRE3 construct to drive expression of h-α-Syn (AAV-h-α-Syn) or empty AAV2/5-CBh-WPRE3 vector containing non-coding (null) stuffer DNA (AAV-EV) into dorsal raphe nucleus (DR) and were euthanized at 4 and 8 weeks (W) post-injection. **A, C, E**) Representative coronal brain image showing SV2A-ir, synaptophysin (SYP)-ir, and MAP-2-ir, respectively in the medial prefrontal cortex (mPFC), cingulate cortex (Cg), caudate-putamen (CPu), and hippocampus (HPC) of mice at different time points after AAV2/5 injection. Scale bar: low: 200 μm. **B, D, F**) Bar chart showing the density of SV2A, SYP, and MAP-2 markers in the brain areas analyzed: prelimbic cortex (PrL), infralimbic cortex (IL), Cg, CPu, and different HPC subfields (n = 5 mice/group; *p< 0.05, **p< 0.01, ***p< 0.001 versus AAV-EV or AAV-h-α-Syn 4W mice). Values are presented as mean ± SEM.

We also examined the density of microtubule-associated protein 2 (MAP-2), a dendritically enriched protein that controls microtubule dynamics, cellular transport, and synaptic function ^54,55^, in AAV-h-α-Syn injected mice compared to control group (**Fig. 2E, 2F**). MAP-2 density was significantly decreased in all brain areas assessed at 8 weeks (reduction of ∼72.3% in prelimbic cortex - PrL, ∼74% in infralimbic cortex - IL, ∼58% in Cg, ∼51% CPu, ∼40% in CA1, and ∼44% in CA2/3; p < 0.001). Taken together, the progressive axonal α-Syn accumulation is associated with an aberrant increase in the density of pre-synaptic markers and a decrease in MAP-2 immunoreactivity throughout the brain, especially after 8 weeks.

We then performed immunoblot analysis on microdissected mouse mPFC, CPu, and HPC extracted 8 weeks after intra-DR injection of AAV-h-α-Syn or AAV-EV. Western-blot (WB) analysis confirmed a regional increase in the levels of the presynaptic markers SV2A (p < 0.01), SYP (p< 0.01), and VAMP2 (vesicle associated membrane protein 2, p< 0.05) in mPFC and marginal effects in CPu (SV2A, p= 0.0507; SYP, p= 0.081; VAMP2, p< 0.05) compared to control group (**Fig. 3A-D**). The changes in synaptic markers in the mPFC, but not in the CPu, were accompanied by a significant increase in the levels of the motor protein dynein (p < 0.01) (**Fig. 3E**). There was also a significant reduction in the post-synaptic marker PSD95 (postsynaptic density protein 95) in the mPFC (p < 0.001) and CPu (p < 0.05) compared to controls (**Fig. 3F**). No changes in these synaptic markers were found in the HPC from mice overexpressing h-α-Syn in DR.

**Figure 3.**
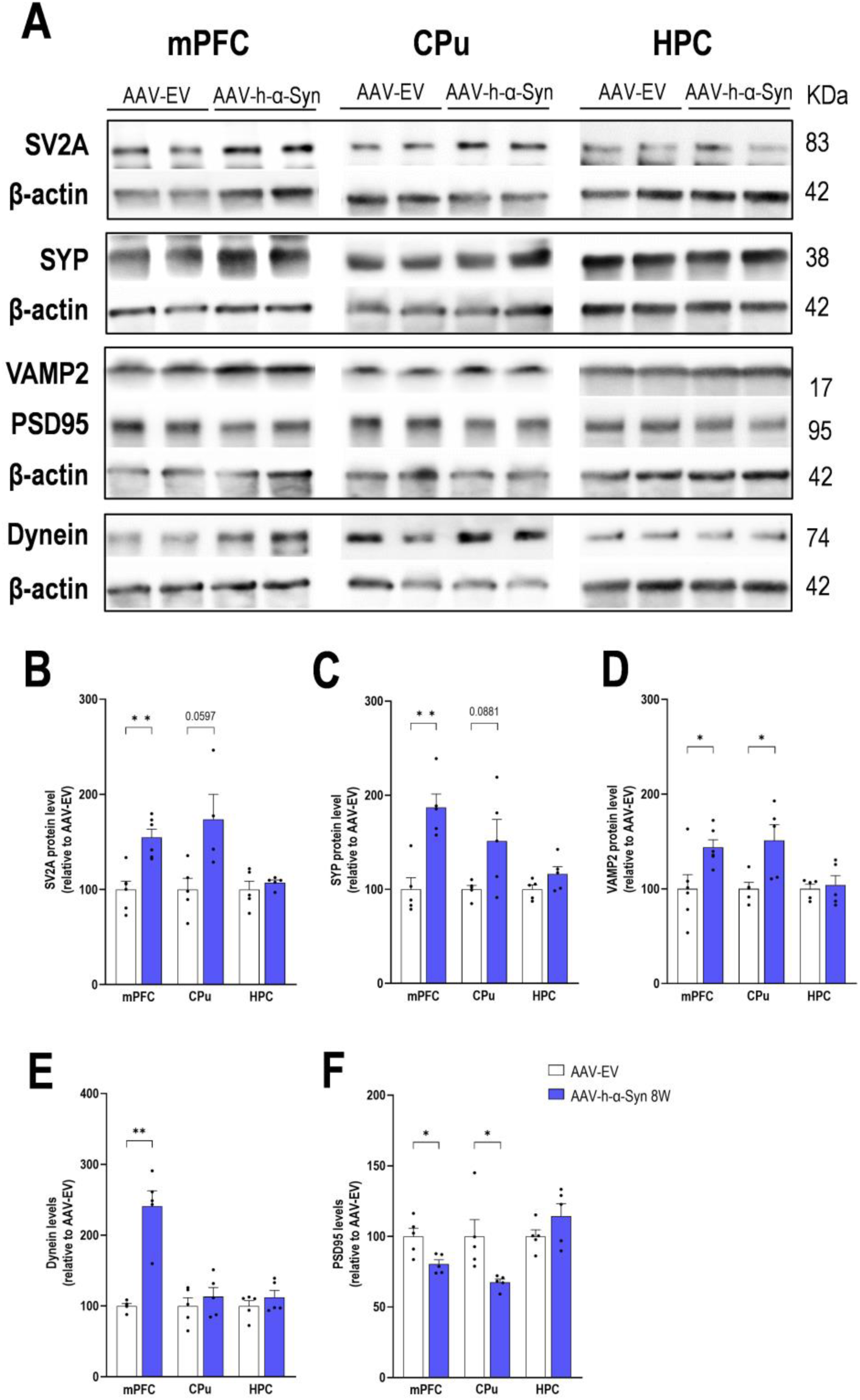
Levels of pre- and post-synaptic and axonal motor proteins in cortical and limbic areas in the D-PD mouse model. Male mice received 1 μl of AAV2/5-CBh-WPRE3 construct to drive expression of h-α-Syn (AAV-h-α-Syn) or empty AAV2/5-CBh-WPRE3 vector containing non-coding (null) stuffer DNA (AAV-EV) into dorsal raphe nucleus (DR) and were euthanized at 8 weeks (W) post-injection. **A)** Western blot of different proteins examined in the medial prefrontal cortex (mPFC), caudate putamen (CPu), and hippocampus (HPC). **B-F)** Bar chart showing the density of SV2A (B), SYP (C), VAMP2 (D), dynein (E), and PSD95 (F) markers in the different brain areas analyzed (n = 5-6 mice/group; *p < 0.05, **p < 0.01, ***p < 0.001 versus AAV-EV). β-actin was used as a protein loading control. Values are presented as mean ± SEM.

### Synaptic pathology in cortical and subcortical brain areas in PD patients

Previously, we showed that the accumulation of oligomeric α-Syn in the dlPFC of PD patients at different Braak stages (early B2-3 and late B5-6) correlates with an increase in PERK-eIF2α signaling, which is known to control the quality of synaptic proteins ^47^. Here, we investigated whether the increased α-Syn levels in different brain areas is associated with regional changes in synaptic markers, as assessed in D-PD-like mice. First, we confirmed the regional presence of oligomeric α-Syn in the dlPFC, CAU, and HPC of PD patients at B2-3 and B5-6 by WB compared to controls (p < 0.01; **Fig. 4A, 4B**). We also detected a significant reduction in SERT levels in the dlPFC and CAU (marginal effects in HPC) examined in both B2-3 and B5-6 stages compared to control group (p < 0.01; **Fig. 4C, 4D**). In parallel, presynaptic SV2A and SYP protein levels were significantly increased in the dlPFC and CAU of PD patients at B2-3 compared to the control group (p < 0.01), but their density decreased in all brain areas examined at B5-6 compared to the control group (p < 0.01; **Fig. 4E-4G**). For presynaptic VAMP2 protein, WB analysis showed higher levels in the dlPFC and CAU at the onset of the disease (B2-3) and decreased levels in CAU and HPC in the late stage (B5-6) compared to the respective control groups (p < 0.01; **Fig. 4E, 4H**). WB analysis of dynein showed marginal increases in motor protein levels in dlPFC (p = 0.0551) and CAU (p = 0.0643) at B2-3 and a slight reduction in HPC (p = 0.0607) at B5-6 compared to controls (**Fig. 4E, 4I**). Furthermore, regionally increased α-Syn levels were associated with reduced PSD95 levels in dlPFC, CAU, and HPC of PD patients at B5-6 compared to the control subjects (p < 0.01; **Fig. 4E, 4J**). Taken together, we found changes in different pre-and post-synaptic markers in cortical and sub-cortical regions that depend on the duration of the disease, with increases in the initial pre-motor phases (B2-3) that progress to decreases in the later stages (B5-6) of the disease, perhaps correlating with cortical/subcortical synaptic loss.

**Figure 4.**
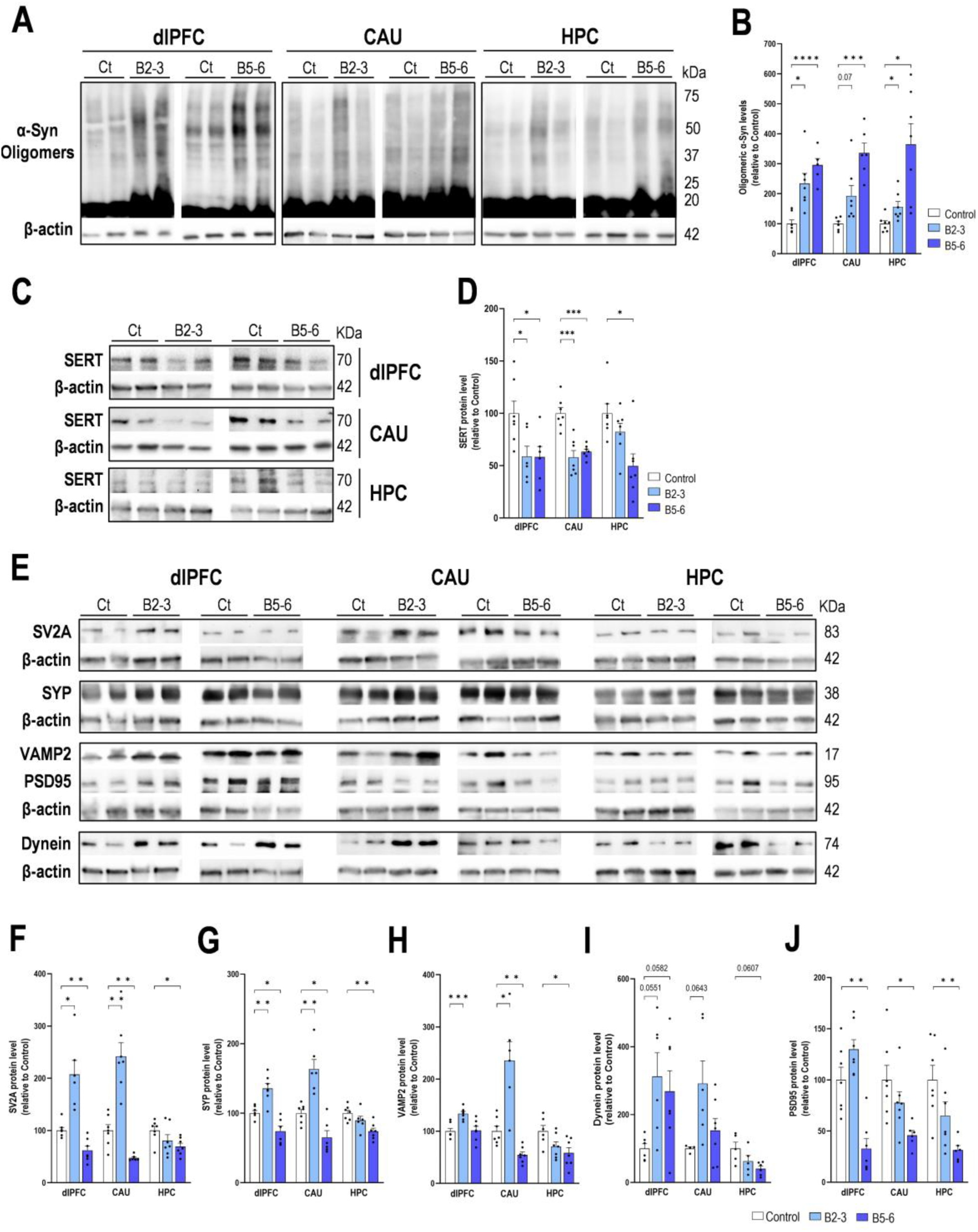
Levels of oligomeric α-Syn, serotonin transporter, synaptic and axonal motor proteins in cortical and limbic areas in early and late PD. **A, B)** Western blot of oligomeric α-Syn in the postmortem samples of dorsolateral prefrontal cortex (dlPFC), caudate nucleus (CAU), and hippocampus (HPC) of PD patients at early and late Braak stages (B2-3 and B5-6, respectively) stages and controls (Ct). Significant increases in the levels of α-Syn oligomers were detected in the dlPFC, CAU, and HPC in PD patients at different stages compared to controls. **C, D)** Western blot of SERT protein in the dlPFC, CAU, HPC in different PD stages and controls. Significant reductions in the SERT levels were detected in the dlPFC, CAU, and HPC in PD patients at different stages compared to controls. **E)** Western blot of different synaptic and motor proteins examined in dlPFC, CAU, and HPC in PD patients and controls. **F-J)** Bar chart showing the density of SV2A (F), SYP (G), VAMP2 (H), dynein (I), and PSD95 (J) markers in the different brain areas analyzed (n = 8 tissue samples/group; *p < 0.05, **p < 0.01, ***p < 0.001 versus controls). β-actin was used as a protein loading control. Values are presented as mean ± SEM.

### Mapping of Egr-1-dependent brain cell activity and functional connectivity in the D-PD-like mouse model

We then investigated whether regional changes in synaptic plasticity translate into changes in Egr-1-dependent whole-brain cellular activity in the D-PD mouse model. We found that Egr-1 mRNA expression assessed by *in situ* hybridization was generally high in the PrL/IL cortices, Cg, CPu, hippocampal area (CA1 and CA2/3), and amygdala (Amg), while expression was lower in the hypothalamus (Hyp) and the DR in AAV-h-α-Syn mice compared to the control group (p < 0.001; **Fig. 5A-C)**. Increases in Egr-1 mRNA expression in efferent cortical and subcortical areas occurred only 8 weeks later, parallel to the changes observed in the synaptic marker levels and the progressive increase in α-Syn-positive fibers. However, the level of Egr-1 mRNA expression was reduced in Hyp and DR 4 weeks later and remained so until the end of the experiment (p < 0.001; **Fig. 5C**).

**Figure 5.**
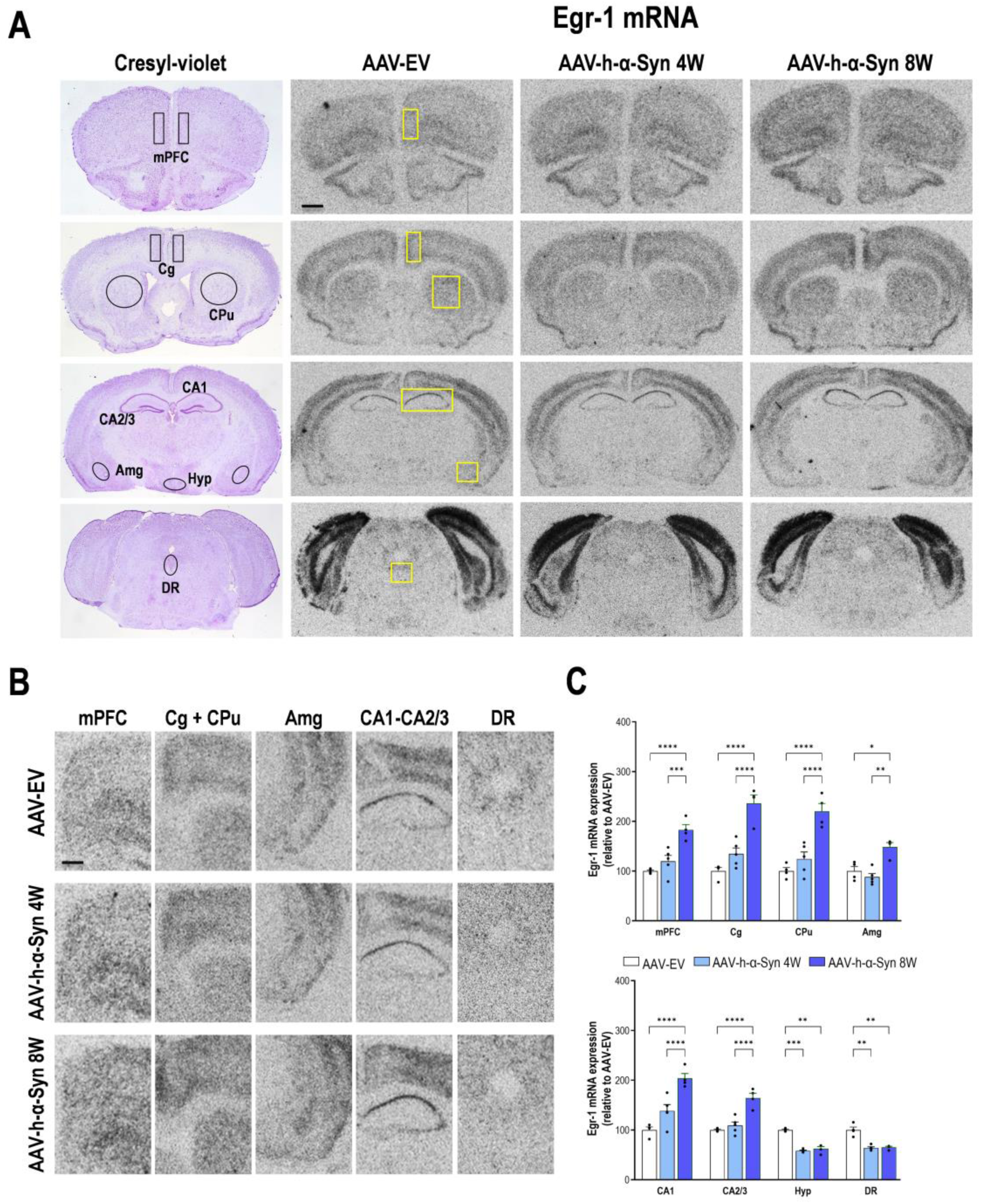
Brain mapping of Egr-1-dependent activity in D-PD mice. Male mice received 1 μl of AAV2/5-CBh-WPRE3 construct to drive expression of h-α-Syn (AAV-h-α-Syn) or empty AAV2/5-CBh-WPRE3 vector containing non-coding (null) stuffer DNA (AAV-EV) into dorsal raphe nucleus (DR) and were euthanized at 4 and 8 weeks (W) post-injection. **A)** Representative images of coronal sections of the mouse brain showing Egr1 mRNA expression in AAV-EV and AAV-h-α-Syn mice assessed by *in situ* hybridization. Scale bar: 1 mm. On the left, cresyl violet-stained coronal sections of the mouse brain show quantified regions of interest (ROI) in the medial prefrontal cortex (mPFC), cingulate cortex (Cg), caudate-putamen (CPu), hippocampal subfields (CA1, CA2/3), amygdala (Amg), hypothalamus (Hyp), and DR. **B)** Enlarged images of the sections of panel A marked with yellow boxes. Scale bar: 100 μm. **C)** Bar chart showing the density of Egr1 mRNA in the different brain areas analyzed (n = 5 mice/group; *p< 0.05, **p< 0.01, ***p< 0.001 versus AAV-EV or AAV-h-α-Syn 4W mice). Values are presented as mean ± SEM.

Given the changed profile of cellular activity in the different brain regions examined, we then characterized brain connectivity in the D-PD-like mouse model at 8 weeks only. We investigated how the net activity of neural elements in a target region is influenced by the net activity of neural elements in a source region using resting-state fMRI (rs-fMRI). In a more specific manner, we evaluated the functional connectivity using seed-based analyses with three different regions considered as seed, namely mPFC, CPu, and HPC. We identified clusters where connectivity was significantly different between groups, and evaluated average connectivity in such areas. In this way, we measured the correlation between brain activity in the regions of interest (seed), automatically identified based on a mouse brain atlas and the rest of the brain. In addition, we analyzed phenotype differences both voxel-wise (**Fig. 6A-C, 7A-C, 8A-C**) and by comparing the average correlation value within each cluster (**Fig. 6D, 7D, 8D**).Voxel-wise statistics with FWE-correction and TFCE did not show differences in the connectivity of mPFC. However, when a p < 0.005 (without FWE-correction) was considered, four cluster of differences were detected.

**Figure 6.**
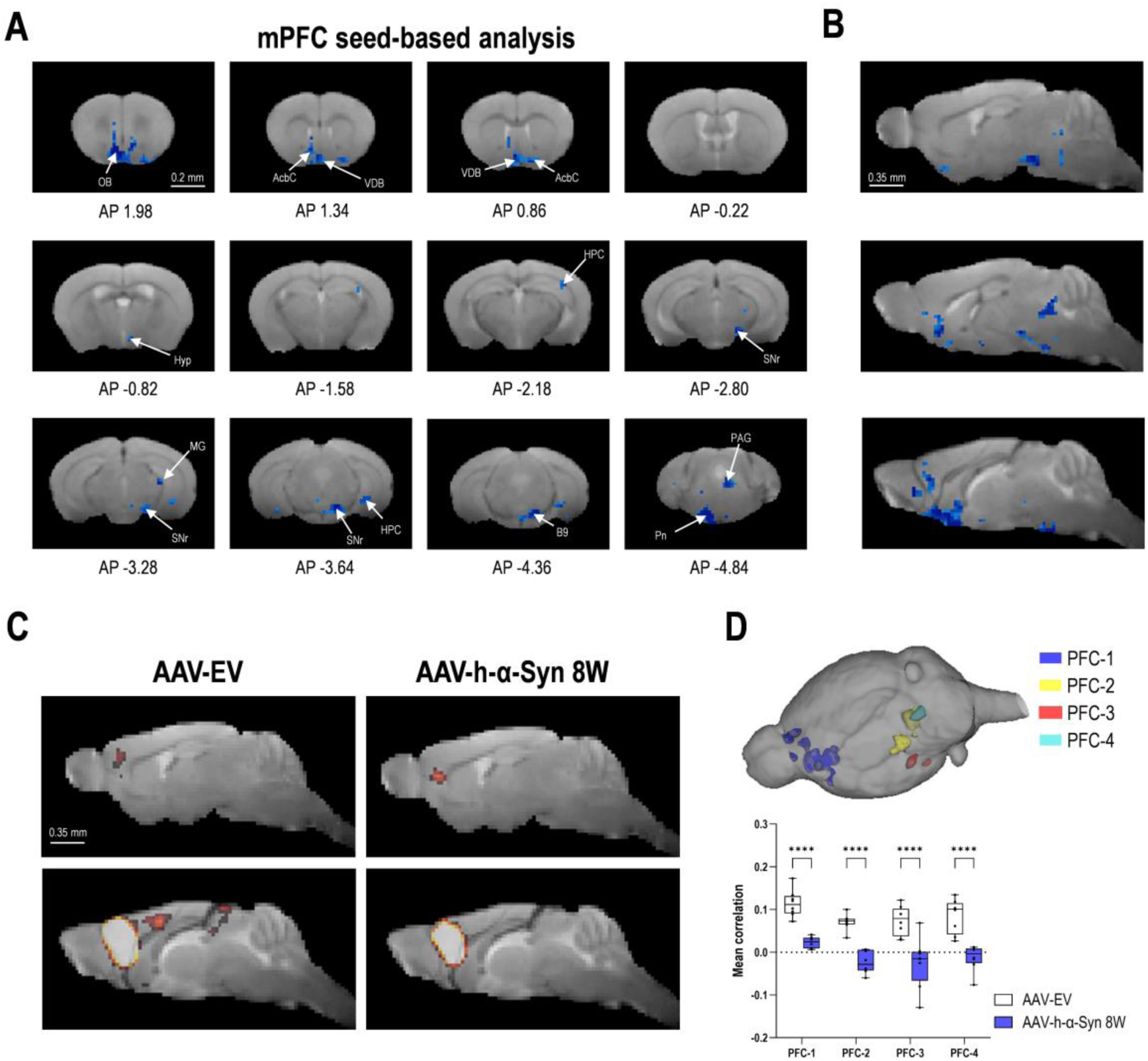
Overexpressing of h-α-Syn in raphe 5-HT neurons in mice results in rsfMRI hypoconnectivity. Seed-based analysis from medial prefrontal cortex (mPFC). Male mice received 1 μl of AAV2/5-CBh-WPRE3 construct to boost h-α-Syn expression (AAV-h-α-Syn) or empty AAV2/5-CBh-WPRE3 vector containing non-coding (null) fill DNA (AAV-EV) in the dorsal raphe nucleus of the dorsal raphe (DR) and were analyzed 8 weeks later. **A, B)** Map of significant differences identified by voxel-wise statistical analysis (randomize with threshold-free cluster enhancement). In blue areas where a decreased connectivity was observed in the AAV-h-α-Syn compared to AAV-EV-injected mice (dark blue, p < 0.005; light blue, p < 0.01). AP: antero-posterior from Bregma in mm. **C)** Average functional connectivity of the mPFC with the rest of the brain. Sagittal slices from the left hemisphere are represented. **D)** For each cluster of voxels with significantly decreased connectivity in the AAV-h-α-Syn compared to AAV-EV-injected mice, average functional connectivity with mPFC is computed as the average of the seed-based correlation map in the specific area. Each gray point represents data from an individual mouse (n = 8 mice/group; *p< 0.05, **p< 0.01, ***p < 0.001 versus controls). Values are presented as mean ± SEM. Abbreviations: AcbC, accumbens nucleus core; B9, serotonin cells; HPC, hippocampus; Hyp, hypothamus; MG, medial geniculate nucleus; OB, olfactory bulb; PAG, periaqueductal gray; Pn, pontine nucleus; SNr, substantia nigra reticulata; VDB, nucleus of the vertical limb of the diagonal band

**Figure 7.**
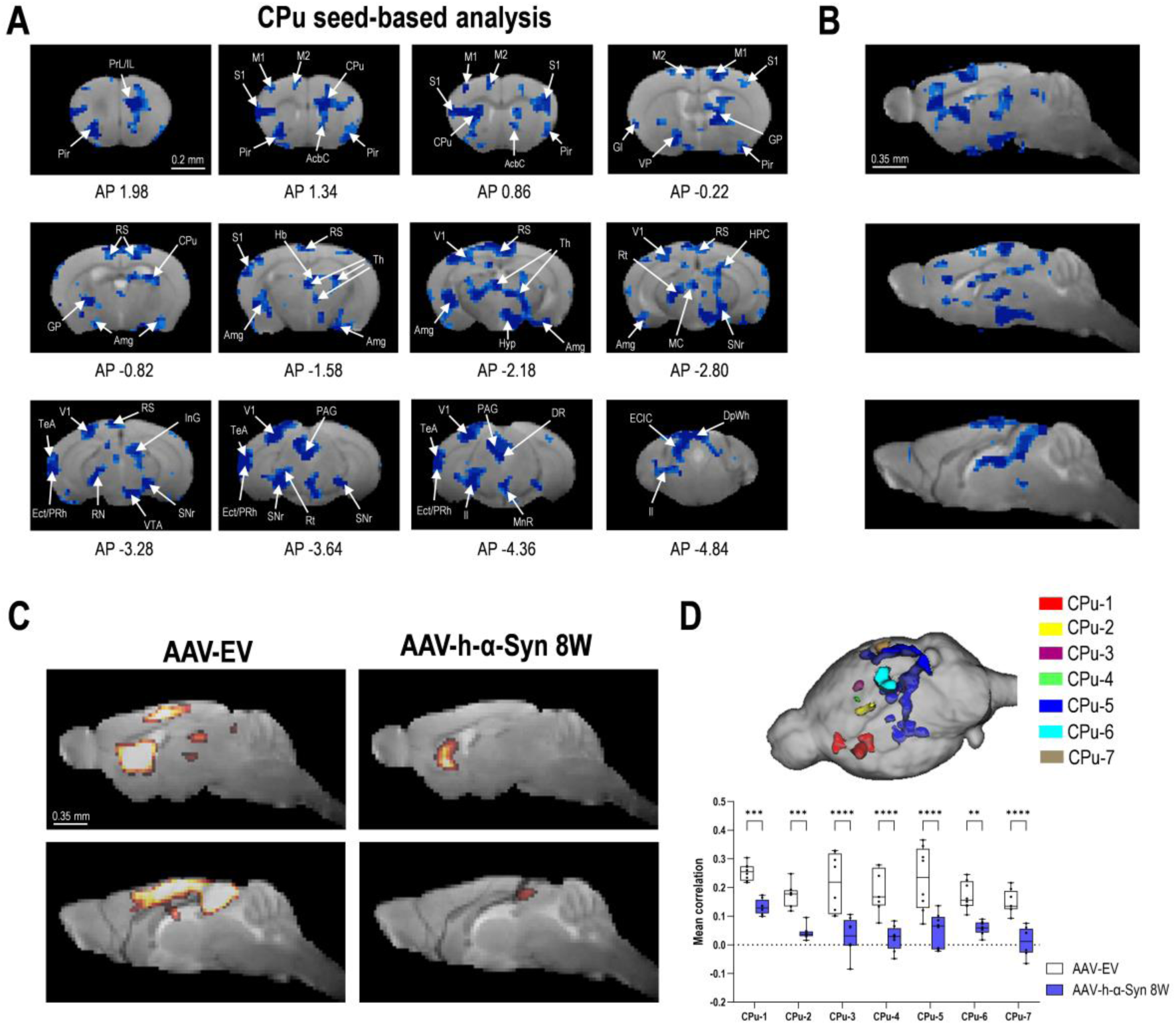
Overexpressing of h-α-Syn in raphe 5-HT neurons in mice results in rsfMRI hypoconnectivity. Seed-based analysis from caudate-putamen (CPu). Male mice received 1 μl of AAV2/5-CBh-WPRE3 construct to boost h-α-Syn expression (AAV-h-α-Syn) or empty AAV2/5-CBh-WPRE3 vector containing non-coding (null) fill DNA (AAV-EV) in the dorsal raphe nucleus of the dorsal raphe (DR) and were analyzed 8 weeks later. **A, B)** Map of significant differences identified by voxel-wise statistical analysis (randomize FWE-correction and threshold-free cluster enhancement). In blue areas where a decreased connectivity was observed in the AAV-h-α-Syn compared to AAV-EV-injected mice (dark blue, corrected p < 0.05; light blue, corrected p < 0.1). AP: antero-posterior from Bregma in mm. **C)** Average functional connectivity of the CPu with the rest of the brain. Sagittal slices from the left hemisphere are represented. **D)** For each cluster of voxels with significantly decreased connectivity in the AAV-h-α-Syn compared to AAV-EV-injected mice, average functional connectivity with CPu is computed as the average of the seed-based correlation map in the specific area. Each gray point represents data from an individual mouse (n = 8 mice/group; *p< 0.05, **p< 0.01, ***p< 0.001 versus controls). Values are presented as mean ± SEM. Abbreviations: AcbC, accumbens nucleus core; Amg, amygdala; CPu, caudate putamen; DpWh, deep white superior coll; DR, dorsal raphe nucleus; ECIC, external cx inferior coll; Ect, ectorhinal cortex; GI, granular insular cx; GP, globus pallidus; HPC, hippocampus; Hyp, hypothamus; IL, infralimbic cortex; InG, intermed gray layer SC; ll, lateral lemniscus; M1/2, primary and secondary motor cortex; MC, magnocellular nucleus; MrR, medial raphe nucleus; PAG, periaqueductal gray; Pir, piriform cortex; PrL, prelimbic cortex; PRh, perirhinal cortex; RS; retrosplenial cortex; Rt, reticular nucleus; S1, primary somatosensorial cortex; SNr, substantia nigra reticulate; TeA, temporal association cortex; Th, thalamus; V1, primary visual cortex; VTA, ventral tegmental area.

**Figure 8.**
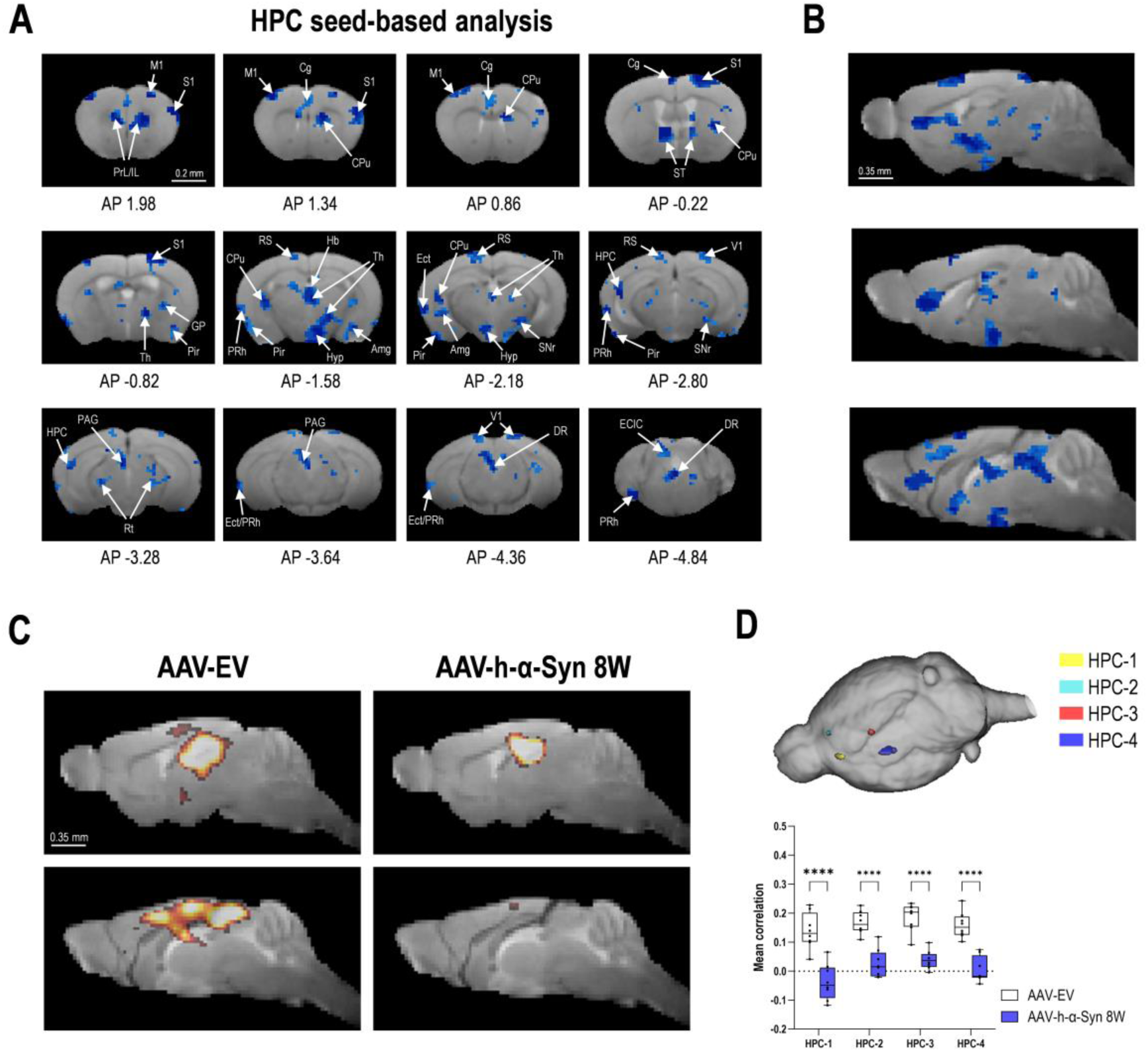
Overexpressing of h-α-Syn in raphe 5-HT neurons in mice results in rsfMRI hypoconnectivity. Seed-based analysis form hippocampus (HPC). Male mice received 1 μl of AAV2/5-CBh-WPRE3 construct to boost h-α-Syn expression (AAV-h-α-Syn) or empty AAV2/5-CBh-WPRE3 vector containing non-coding (null) fill DNA (AAV-EV) in the dorsal raphe nucleus of the dorsal raphe (DR) and were analyzed 8 weeks later. A, B) Map of significant differences identified by voxel-wise statistical analysis (randomize FWE-correction and threshold-free cluster enhancement). In blue areas where a decreased connectivity was observed in the AAV-h-α-Syn compared to AAV-EV-injected mice (dark blue, corrected p<0.05; light blue, corrected p<0.1). AP: antero-posterior from Bregma in mm. C) Average functional connectivity of the HPC with the rest of the brain. Sagital slices from the left hemisphere are represented. D) For each cluster of voxels with significantly decreased connectivity in the AAV-h-α-Syn compared to AAV-EV-injected mice, average functional connectivity with HPC is computed as the average of the seed-based correlation map in the specific area. Each gray point represents data from an individual mouse (n = 8 mice/group; *p< 0.05, **p< 0.01, ***p< 0.001 versus controls). Values are presented as mean ± SEM. Abbreviations: Amg, amygdala; CPu, caudate putamen; Cg, cingulate cortex; DR, dorsal raphe nucleus; ECIC, external cx inferior coll; Ect, ectorhinal cortex; GP, globus pallidus; Hb, habenular commissure; Hyp, hypothamus; IL, infralimbic cortex; M1, primary motor cortex; Pir, piriform cortex; PrL, prelimbic cortex; PRh, perirhinal cortex; RS; retrosplenial cortex; Rt, reticular nucleus; S1, primary somatosensorial cortex; SNr, substantia nigra reticulate; ST, bed nucleus of stria terminalis; Th, thalamus; V1, primary visual cortex.

Compared to control group, the left mPFC of AAV-h-α-Syn mice showed reduced functional connectivity with the cluster 1 to 4 named as: PFC-1 (medial orbital cortex, PrL/IL, and accumbens nucleus; p < 0.0001), PFC-2 (substantia nigra reticulata, pontine reticular nucleus, and B9 5-HT cells; p < 0.0001), PFC-3 (HPC; p < 0.0001), and PFC-4 (DR and periaqueductal gray; p < 0.0001) (**Fig. 6A-D**).

Furthermore, we also evaluated functional connectivity through a seed-based analysis between the left CPu and the rest of the brain (**Fig. 7A-D**). Compared to control group, the left CPu of AAV-h-α-Syn mice showed reduced functional connectivity with clusters CPu-1 to CPu-7 (corrected p-value < 0.05): CPu-1 (Prl/IL, CPu, somatosensory cortex, and globus palillus; p < 0.01), CPu-2 (motor cortex 1/2; p < 0.0001), CPu-3 and CPu-6 (retrosplenial cortex; p < 0.0001), CPu-5 (habenula, thalamus, hypothalamus, HPC, amygdala, substantia nigra reticulata, ventral tegmental area, visual cortex, DR, and periaqueductal gray; p < 0.0001), CPu-4 (CPu; p < 0.0001), and CPu-7 (ectorhinal and perirhinal cortices; p < 0.01).

Finally, we also evaluated functional connectivity through a seed-based analysis between the left HPC and the rest of the brain (**Fig. 8A-8C**). Compared to control group, the left HPC of AAV-h-α-Syn mice showed reduced functional connectivity with clusters HPC-1 to HPC-7 (corrected p-value < 0.05): HPC-1 (medial orbital cortex, and PrL/IL cortices; p < 0.0001), HPC-2 (CPu; p < 0.0001), HPC-3 (somatosensory cortex; p < 0.0001), and HPC-4 (thalamus and hypothalamus; p < 0.0001). Overall, our results highlight that cortico-limbic-raphe functional connectivity is greatly affected in D-PD-like mice.

## DISCUSSION

Depression is one of the most common non-motor symptoms, is highly prevalent in PD patients, and one of the most important predictors of poor quality of life in people with PD. It often precedes motor symptoms, but is underdiagnosed and undertreated ^56,57^. Indeed, the diagnosis of depression in PD is complex, especially when different clinical presentations of depression and PD overlap. In recent years, neuroimaging reports have suggested reduced connectivity between the cortico-limbic networks, perhaps as a result of altered higher-order cortical modulatory effects in emotion-related limbic areas (including the thalamus, ventral striatum, insula, Amg, and Cg), leading to mood dysregulation ^33,37,58,59^. In support of this, multimodal neuroimaging studies combining structural and connectivity analysis have been able to identify neurophysiological subtypes of PD patients, distinguishing between those with the dominant severe depression subtype and those with the dominant severe movement subtype. The first group have an altered connectivity pattern, mainly composed in limbic regions ^56^. An important biological basis for depression in PD is the result of the impairment of 5-HT innervation and neurotransmission in brain areas of the cortico-limbic system ^6,14,15,17,60^. We have previously developed a mouse model with overexpression of h-α-Syn in DR 5-HT neurons that exhibits depressive-like behavior associated with reduced 5-HT neurotransmission in the mPFC and CPu ^45^. Here, we extended these studies and showed that axonal h-α-Syn accumulation in mice overexpressing the transgene in DR is coupled to a progressive loss of 5-HT SERT-positive fibers in cortical and subcortical areas and to a functional hypo-connectivity in cortico-limbic-raphe network, mainly at 8 weeks. Using seed-based analyses of rs-fMRI, we identified major clusters of connectivity changes between the mPFC-Accumbes nucleus, mPFC-DR, CPu-PrL/IL, CPu-thalamus + Amg, CPu-HPC, HPC-PrL/IL, HPC-CPu, and HPC-thalamus, networks which may be specifically involved in different aspects of emotion behavioral regulation, as reported in patients with D-PD. It is important to highlight the hypo-connectivity found between the mPFC-DR and CPu-DR using this mouse model of D-PD. Previous neuroimaging studies have reported a unique profile of reduced connectivity between the DR and the PFC + Cg that correlated with anxiety, drowsiness, and depression in PD ^61^. Taken together, the data support the translational nature of this mouse model for furthering our understanding of the neurobiological substrates underlying depression in PD.

In parallel with the changes in the activity of the cortico-limbic-raphe network, we found an increase in Egr-1-dependent cellular activity in the cortico-limbic regions (PrL/IL, Cg, CPu, HPC, and Amg) in the D-PD mouse model at 8 weeks. Egr-1 (also called NGFI-A, Zif268, or Krox24) is a zinc finger transcription factor implicated in processes of synaptic plasticity and cell activation, and has been shown to have a crucial role in both neuronal death and inflammatory response ^62–64^. In fact, elevated Egr-1 levels were detected in a PD mouse model of 1-methyl-4-phenyl-1,2,3,6-tetrahydropyridine (MPTP) associated with an astrocyte activation and neuroinflammation ^64^. Although we have not identified the Egr-1-dependent activated cell type in the whole brain, we have found increases in the neuroinflammatory markers TNFα, INFγ, and NFkB1 in the mPFC and CPu using this D-PD mouse model, suggesting a possible Egr1-dependent activation mechanism leading to astrogliosis and/or microgliosis (Miquel-Rio et al., personal communication). In addition, we also found reduced Egr-1 expression in the DR of D-PD mice locally injected with AAV-h-α-Syn. DR –one of the most extensively connected hubs in the mammalian brain– is the largest 5-HT nucleus, containing approximately one third of all 5-HT neurons in the brain ^49^, although clusters of GABAergic, DAergic, and to a lesser extent, glutamatergic and peptidergic neurons have been identified in the DR ^65^. In the present study, no changes in the number of TPH-positive or TH-positive cells were detected in the DR after h-α-Syn accumulation, but a reduced density of the SERT-positive fibers was shown in cortico-limbic regions, suggesting changes in the 5-HT system neuroplasticity. This could translate into a reduction in the specific activity of 5-HT cells, as observed with a reduction in Egr-1 mRNA expression in the DR. These results are further supported by our previous observations, confirming that overexpression of h-α-Syn in the mouse DR leads to decreased 5-HT neuron activity and 5-HT release in mPFC and CPu ^45^.

As mentioned above, recent PET neuroimaging studies have demonstrated specific 5-HT denervation in cortico-striatal-limbic areas in PD patients with prominent neuropsychiatric signs at disease onset ^15,18^. Although some TH-positive fibers were also positive for h-α-Syn in projection brain areas (e.g. Cg and CPu) after AAV-h-α-Syn infusion in the DR, a large proportion of h-α-Syn-positive fibers colocalized with SERT-positive 5-HT fibers. More importantly, a reduction in SERT-positive fibers density, but not TH-positive fibers, was detected in the cortico-striatal limbic regions (CPu > Cg > mPFC > HPC) in D-PD mice compared to controls, as previously shown in animal models of h-α-Syn overexpression in non-DA neurons ^66,67^. It is known that α-Syn is a protein that regulates the expression and cellular trafficking of monoamine transporters, e.g. SERT, DA transporter – DAT, and norepinephrine transporter - NET, towards the cell surface ^16^. α-Syn-induced modulation of SERT trafficking is microtubule-dependent, as the microtubule destabilizing agent nocodazole disrupts the effects of α-Syn on SERT function and reverses the inhibition of uptake in co-transfected cells ^68^. In addition, *in vivo* studies have shown that changes in the level of α-Syn expression in mouse 5-HT neurons directly produce opposite effects on the functional activity of SERT in efferent brain regions in response to the inhibitor citalopram ^45,69^. Taken together, this may suggest that the progressive accumulation of h-α-Syn in 5-HT axons and terminals leads to impaired SERT function in serotonergic synapses, resulting in the loss of system integrity.

Actually, synaptic dysfunction and altered axonal transport have been postulated to be some of the earliest pathological events in PD, in which α-Syn is considered a hub protein by interacting with many synaptic proteins (e.g. monoamine transporters), cytoskeletal components, chaperones, and several synaptic vesicles SV-associated proteins ^52,70,71^. In physiological conditions, α-Syn is abundant in the pre-synaptic compartment ^51^. Post-mortem and experimental studies provided evidence that α-Syn aggregation might start at pre-synaptic terminals, inducing synaptic degeneration and subsequently neuronal loss ^52,70^. Unlike previous studies that showed a decrease in the density of pre-synaptic markers SV2A, SYP, and VAMP2 in aged PD mice (17-20 months) ^72–75^, here we reported an accumulation of these same pre-synaptic proteins in parallel with higher h-α-Syn levels, mainly in the Cg and CPu in the D-PD mouse model. Indeed, we have previously shown that increased levels of SV2A protein co-localize with h-α-Syn in axonal swellings in mouse CPu and Cg ^45^. In this regard, increases of the presynaptic SV2C protein - enriched in the basal ganglia and preferentially localized in DA neurons-were also reported in mice overexpressing mutant A53T α-Syn ^76,77^. It is important to emphasize that we also observed increases in the levels of SV2A, SYP and VAMP2 proteins in the dlPFC and CAU of patients with PD in the early stages B2-3 of the disease, but these decreased in the later stages B5-6. Our data are consistent with recent studies showing reduced SYP and SV2A densities in cortical regions in B5-6 α-Syn stage compared to controls, associated with higher neurofilament light-chain (NfL) immunoreactivity and Lewy body density ^71,78^. Although one of the main limitations of our study is the small number of post-mortem human tissue samples and the absence of an accurate diagnosis of depressive disorder in PD patients, the data support the idea that synaptic degeneration can occur together with axonal degeneration in the final stage of the disease. In the early B2-3 stages, when non-motor symptoms prevail, changes in the trafficking/refilling of synaptic vesicles mediated by α-Syn and/or in the structural organization of synapse may contribute to functional deficits in different brain networks, including those that regulate emotional processes. In fact, our data and others support these observations by showing reductions in the density of the 5-HT SERT marker that directly interacts with α-Syn in both stages B2-3 and B5-6 of PD ^79,80^.

In this study, we also found a significant increase in the levels of axonal transport motor protein dynein in the mPFC in the D-PD mouse model overexpressing h-α-Syn in DR. Dynein is a microtubule-based motor protein that mediates minus-end-directed vesicle transport through interaction with dynactin ^55,81^. Previous studies have reported that mice and rats overexpressing A53T α-Syn have higher dynein levels compared to the control group, leading to an accumulation of pre-synaptic proteins and a disruption of axonal transport, which precedes axonal denervation ^82,83^. We observed a similar pattern here, with a cortical loss of 5-HT fibers associated with increased levels of h-α-Syn and vesicular proteins. In addition, reductions in dynein levels were reported in substantia nigra and putamen at late PD stages ^84^. Although we detected marginal effects in specific brain regions, these were increases in early stages B2-3 of PD (e.g. dlPFC and CAU), followed by decreases in stages B5-6 of PD (e.g. HPC), supporting that neurodegeneration in PD involves a breakdown of axonal transport. Remarkably, the changes in pre-synaptic plasticity that occurred in the D-PD mouse model led to changes at the post-synaptic level, with decreased MAP-2 (localized primarily in neuronal dendrites) and PSD95 levels in brain areas of the cortico-limbic network that could be linked to the detected functional hypo-connectivity. Like other authors, we also detected reduced levels of PSP95 in advanced stages of PD (Esteves and Cardoso, 2020; but not Fourie et al., 2014; Whitfield et al., 2014)

In conclusion, we have shown here that overexpression of h-α-Syn in the 5-HT system leaves to a hypo-connectivity in the cortico-limbic-raphe network that closely resembles the functional changes described in patients with depression and PD. Elevated levels of h-α-Syn in 5-HT projection areas trigger an abnormal pattern of accumulation of synaptic vesicle-associated proteins and defects in dynein-driven transport that would lead to reduced 5-HT neurotransmission, alterations in postsynaptic plasticity in efferent brain areas and the development of a depressive phenotype. These findings may help to better understand the mechanisms underlying depression in PD. This D-PD mouse is a powerful translational model for studying new targets and potential antidepressant interventions in α-Syn-associated neurodegenerative disorders.

## METHODS

### Mice

Male and female C57BL/6J mice (10 weeks; *n=*254 for the whole study, Charles River, Lyon, France) were housed under controlled conditions (22 ± 1°C; 12h light/dark cycle) with food and water available ad-libitum. Animal procedures were conducted in accordance with standard ethical guidelines (EU directive 2010/63 of 22 September 2010) and approved by the local ethical committee (University of Barcelona).

### Mouse model overexpressing h-α-Syn into serotonin (5-HT) neurons

Recombinant adeno-associated viral vector serotype-2/5 (AAV2/5) with chicken-β-actin (CBh) promoter and a shortened WPRE3 that encodes the human wild-type α-synuclein (AAV-h-α-Syn) was used to overexpress the transgene in raphe 5-HT neurons of mice (dorsal raphe nucleus - DR), as previously reported ^45^. The Michael J. Fox Foundation gently provided the AAV2/5 constructs produced and tittered by the UNC Vector Core. We examined any possible effect of the AAV2/5 vector itself capable of exerting toxicity on 5-HT neurons and causing behavioral alterations using an empty AAV2/5-CBh-WPRE3 vector containing non-coding (null) stuffer DNA (AAV-EV) as a control group. Isoflurane-anesthetized mice (doses: 4% induction, 2% maintenance) were randomly injected with 1 μl of AAV-h-α-Syn (concentration 0.5 x 10^13^ gc/ml) or AAV-EV (concentration 0.5 x 10^13^ gc/mL) into raphe nuclei (anterior-posterior AP: -4.5; medial-lateral ML: -1.0; dorsal-ventral DV: -3.2 in mm, relative to bregma with an angle 20°), using a microinjector (KDS-310-PLUS, World Precision Instruments, Sarasota, LF, USA) at 0.4 µl/min rate. The needle was kept in place for an additional 5 min before slowly being withdrawn. Mice were assessed at 4 and 8 weeks after AAV-h-α-Syn or AAV-EV infusion.

### Postmortem human brain samples

Dorsolateral prefrontal cortex (dlPFC), caudate nucleus (CAU), and hippocampus (HPC) tissue samples from controls and PD patients were obtained from the Hospital Clinic-IDIBAPS biobank, in accordance with Spanish legislation and with the approval of the Ethics Committee of Spanish National Research Council (Project Ref. PID2019-105136RB-I00). The post-mortem interval between death and the processing of the tissues ranged from 3 to 20 h, and the samples were stored at -80°C until they were tested. Pathological cases were categorized as stages 1-6 of LB disease pathology, according to Braak staining ^12^, excluding tauopathies, vascular disease, and metabolic syndrome. No information (e.g. by formal testing) could be obtained on the neuropsychiatric status of the PD patients. Cases in the control group had not neurological, psychiatric, or metabolic disorders and no neuropathological abnormalities, except for sporadic Alzheimer’s disease. A total of 8 control and 16 cases with PD-related pathology were included in the present study. All PD cases were treated for motor symptoms. The duration of the disease was in the range of 8 to 18 years. The most common causes of death in people with PD were infections, neoplasia, and acute heart disease. See **Table 1** for subject details.

**Table 1.** Demographic and clinical characteristics of each group used in the present study.

|  | Number of subjects | Gender |  | Age (year) |  | Postmortem interval (hour) |  |
| --- | --- | --- | --- | --- | --- | --- | --- |
|  |  | Male | Female | Male | Female | Male | Female |
| Control | 8 | 3 | 5 | 74.7 ± 5.5 | 78.0 ± 5.1 | 8.5 ± 1.7 | 15.3 ± 2.1 |
| Braak 2-3 | 8 | 4 | 4 | 77.7 ± 4.9 | 88.0 ± 6.4 | 8.7 ± 1.7 | 9.7 ± 2.3 |
| Braak 6-6 | 8 | 4 | 5 | 78.5 ± 3.5 | 76.2 ± 4.5 | 9.4 ± 3.6 | 9.0 ± 3.0 |
| P value | N/A | N/A | N/A | 0.842 | 0.311 | 0.972 | 0.168 |

Values denote mean ± SEM. PD: Parkinson's disease, N/A: not applicable.

### *In situ* hybridization (ISH)

Mice were euthanized by cervical dislocation and brains were rapidly removed, frozen on dry ice and stored at -80°C. Coronal tissue sections (14 μm-thick) containing medial prefrontal cortex (mPFC), cingulate cortex (Cg), caudate-putamen (CPu), hippocampus (HPC), amygdala (Amg), hypothalamus (Hyp), and dorsal raphe nucleus (DR) were obtained and processed, as described elsewhere ^88,89^. An antisense oligo probe sequence (gcatcatctcctccagtttggggtagttgtccatggtgggtgagt), complementary to bases Egr1 (NM_007913.5) was used (IBA Nucleic Acids Synthesis, Göttingen, Germany). Frozen tissue sections were fixed for 20 min at 4°C in 4% paraformaldehyde in phosphate-buffered saline (1x PBS: 8 mM Na_2_HPO_4_, 1.4 mM KH_2_PO_4_, 136 mM NaCl, and 2.6 mM KCl), washed for 5 min in 3xPBS at room temperature, twice for 5 min each in 1x PBS, and incubated for 2 min at 21°C in a solution of predigested pronase (Merck Millipore, Madrid, Spain) at a final concentration of 24 U/mL in 50 mM Tris-HCl, pH 7.5, and 5 mM EDTA. The enzymatic activity was stopped by immersion for 30 s in 2 mg/ml glycine in 1x PBS. Tissues were finally rinsed in 1xPBS and dehydrated through a graded series of ethanol.

Oligonucleotide was individually labeled (2 pmol) at the 3’-end with [^33^P]-dATP (>2500 Ci/mmol; DuPont-NEN, Boston, MA, USA) using terminal deoxynucleotidyl-transferase (TdT, Calbiochem, La Jolla, CA, USA). For hybridization, the radioactively labelled probes were diluted in a solution containing 50% formamide, 4x standard saline citrate, 1x Denhardt’s solution, 10% dextran sulfate, 1% sarkosyl, 20 mM phosphate buffer, pH 7.0, 250 μg/ml yeast tRNA, and 500 μg/ml salmon sperm DNA. The final concentration of radioactive probes in the hybridization buffer was in the range (∼1.5 nM). Tissue sections were covered with hybridization solution containing the labelled probes, overlaid with parafilm coverslips and incubated overnight at 42°C in humid boxes. Sections were then washed 4 times (45 min each) in a buffer containing 0.6 M NaCl and 10 mM Tris-HCl (pH 7.5) at 60°C. Hybridized sections were exposed to Biomax-MR film (Sigma-Aldrich) for 2-10 days at −70 °C with intensifying screens. For specificity control, adjacent sections were incubated with an excess (50x) of unlabeled probes. Films were analyzed and relative optical densities were evaluated in three adjacent sections by duplicate of each mouse, and averaged to obtain individual values using Image-J (v1.54c, NIH, Bethesda, MD, USA) software. Contrast and brightness of images were the only variables digitally adjusted.

### Immunofluorescence

Mice were anaesthetized with pentobarbital and transcardially perfused with 4% PFA in sodium-phosphate buffer (pH 7.4). Brains were extracted, post-fixed 24 h at 4°C in the same solution, and placed in gradient sucrose solution 10–30% for 3 days at 4°C. After cryopreservation, serial 30 μm-thick sections were cut to obtain raphe nuclei, mPFC, Cg, CPu, and HPC. Immunofluorescence procedure was performed for h-α-Syn (anti-h-α-Synuclein clone Syn 211, 1:2000; ref.: AHB0261, Thermo Fisher Scientific, Waltham, MA, USA), mouse/human-α-Syn (anti-m/h-α-Syn, 1:500; ref.: ab212184, Abcam, Cambridge, UK) phospho-S129-α-Syn (anti-phospho-S129-α-Syn 1:2000; ref.: ab51253, Abcam, Cambridge, UK), tryptophan hydroxylase TPH (anti-TPH, 1:2500; ref.: AB1541, Sigma-Aldrich, Madrid, Spain), serotonin transporter SERT (anti-SERT, 1:2500; ref.: 24330, Immunostar, Hudson, WI, USA), glutamate decarboxylase GAD67 (anti-GAD67, 1:2500; ref.: MAB5406, Sigma-Aldrich, Madrid, Spain), tyrosine hydroxylase TH (anti-TH, 1:1250; ref.: ab112, Abcam, Cambridge, UK), MAP-2 (anti-MAP-2, 1:1000; ref.: ab32454, Abcam, Cambridge, UK), SV2A (anti-SV2A, 1:250; ref.: PA5-110451, Thermo Fisher Scientific, Waltham, MA, USA), and synaptophysin (anti-SYP, 1:1000; ref.: ab32127, Abcam, Cambridge, UK). Sections were washed in 1x PBS (pH 7.4) and 1x PBS/Triton 0.2%, and incubated in blocking solution (1x PBS/Triton containing 0.02% gelatin and the corresponding normal serum for the secondary antibody host) for 2 hours at room temperature. Primary antibody was incubated overnight at 4°C, followed by incubation with the respective secondary antibodies for 2 hours at room temperature (**Table 2**). Finally, nuclei were stained with Hoestch dye (1:10.000; ref.: H3570, Life Technologies, Carlsbad, CA, USA) for 10 min and the sections were mounted in Entellan (Electron Microscopy Sciences, Hatfield, PA, USA).

**Table 2.** List of antibodies used in the study.

| Protein | Antibody | Company. Ref. | Blocking serum | Primary antibody dilution | Secondary antibody dilution |
| --- | --- | --- | --- | --- | --- |
| Immunofluorescence |  |  |  |  |  |
| h-α-Syn | Mouse anti-h-α-Syn | Invitrogen. AHB0261 | Goat 5% | 1:2000,<br>Donkey serum 2% | 1:500, A488<br>Goat anti-mouse |
| m:h-α-Syn | Rabbit anti-m/h-α-Syn | Abcam. ab212184 | Donkey 5% | 1:500,<br>Donkey serum 2% | 1:500, A488<br>Donkey anti-rabbit |
| P-S129-α-Syn | Rabbit anti-α-Syn<br>Phospho S129 | Abcam. ab51253 | Donkey 5% | 1:2000,<br>Donkey serum 2% | 1:500, A488<br>Donkey anti-rabbit |
| SERT | Rabbit anti-SERT | ImmunoStar. 24330 | Donkey 5% | 1:2500,<br>Donkey serum 2% | 1:500, A555<br>Donkey anti-rabbit |
| TPH | Sheep anti-TPH | Millipore. AB1541 | Donkey 5% | 1:2500,<br>Donkey serum 2% | 1:500, A555<br>Donkey anti-sheep |
| TH | Rabbit anti-TH | Abcam. ab112 | Donkey 5% | 1:1250,<br>Donkey serum 2% | 1:500, A555<br>Donkey anti-rabbit |
| GAD67 | Mouse anti-GAD67 | Millipore. MAB5406 | Rabbit 5% | 1:2500,<br>Rabbit serum 2% | 1:500, A555<br>Rabbit anti-mouse |
| MAP2 | Rabbit anti-MAP2 | Abcam. ab32454 | Donkey 5% | 1:1000,<br>Donkey serum 2% | 1:500, A555<br>Donkey anti-rabbit |
| SYP | Rabbit anti-SYP | Abcam. ab32127 | Donkey 5% | 1:1000,<br>Donkey serum 2% | 1:500, A555<br>Donkey anti-rabbit |
| SV2A | Rabbit anti-SV2A | Fisher. PA5-110451 | Donkey 5% | 1:250,<br>Donkey serum 2% | 1:500, A555<br>Donkey anti-rabbit |
| Western Blot |  |  |  |  |  |
| m/h-α-Syn | Rabbit anti-m/h-α-Syn | Abcam. ab212184 | BSA 5% | 1:1000, BSA 5% | 1:25000, HRP<br>Goat anti-rabbit, BSA 5% |
| SERT | Rabbit anti-SERT | Abcam. ab102048 | Milk 5% | 1:1000, BSA 5% | 1:25000, HRP<br>Goat anti-rabbit, Milk 5% |
| SYP | Rabbit anti-SYP | Abcam. ab32127 | Milk 5% | 1:1000, BSA 5% | 1:25000, HRP<br>Goat anti-rabbit, Milk 5% |
| Dynein | Mouse anti-Dynein<br>intermediate chain 1 | Millipore. MAB1618 | Milk 5% | 1:2000, BSA 5% | 1:25000, HRP<br>Goat anti-mouse, Milk 5% |
| VAMP2 | Rabbit anti-VAMP2 | Abcam. ab3347 | Milk 5% | 1:2000, BSA 5% | 1:25000, HRP<br>Goat anti-rabbit, Milk 5% |
| PSD95 | Mouse anti-PSD95 | Abcam. ab2723 | Milk 5% | 1:3000, BSA 5% | 1:25000, HRP<br>Goat anti-mouse, Milk 5% |
| SV2A | Rabbit anti-SV2A | Fisher. PA5-110451 | Milk 5% | 1:1000, BSA 5% | 1:25000, HRP<br>Goat anti-rabbit, Milk 5% |
| B-actin | Mouse anti-β-actin-HRP | Merck. A3854 | BSA 5% | 1:50000, BSA 5% | - |

### Confocal fluorescence microscopy

The intracellular localization of h-α-Syn or p-α-Syn proteins in TPH^+^ or GABA^+^ cells in the DR, and the intracellular h-α-Syn distribution in 5-HT axons were examined by confocal microscopy using an inverted Nikon Eclipse Ti2-E microscope (Nikon Instruments, Tokyo, Japan) attached to the spinning disk unit Andor Dragonfly 200 (Oxford Instruments Company, Abindong, UK). For all experiments, a Plan Apochromatic 10-20x, numerical aperture (NA) 0.45 objective was used. A high-precision motorized stage was used to collect the large-scale 3D mosaics of each tissue section. Individual image tiles were 2048 × 2048 pixels with a z-section of 35 µm. For specific regions, we used an oil-immersion objective (Plan Apochromatic Lambda blue 40x, NA 1.4). Samples were excited with 405, 488, and 561 nm laser diodes, respectively. The beam was coupled into a multimode fiber going through the Andor Borealis unit reshaping the beam from a Gaussian profile to a homogenous flat top, and from there it was passed through the 40 µm pinhole disk. Tissue sections were imaged on a high resolution scientific complementary metal oxide semiconductor (sCMOS) camera (Zyla 4.2, 2.0 Andor, Oxford Instruments Company). Fusion software (Andor, Oxford Instruments Company) was used for acquisition and for image processing before analysis. Image stitching and deconvolution were performed using Fusion software (Andor, Oxford Instruments Company). Image analysis was performed with Image J/Fiji open source software (Wayne Rasband, NIH, Bethesda, MD, USA) using ImageJ Macro Language to develop custom Macros. Briefly, maximum projections of the channels of interest (red and green) were obtained for each section. Region of interest (ROI) was delimited for each brain section so that the maximum area could be measured in all the images acquired.

Various measurements were applied to the resulting images. For images co-labelling m/h- α-Syn/TPH or h-α-Syn/TH in the DR, the number of cells in the red channel was quantified after adjusting and homogenizing pixel values. Red-labeled cells were then examined for colocalization with the green channel upon merging and subsequently counted. Due to insufficient contrast with the background, GAD67+ cells could not be directly counted. Instead, for images co-labelling p-α-Syn/GAD67 in the DR, total p-α-Syn+ cells were quantified, and then those colocalizing with GAD67 were identified. In projection brain areas (mPFC, Cg, CPu, HPC), images co-labelling h-α-Syn/SERT and h-α-Syn/TH were acquired, and the mean intensity values for the green and red channels were measured for each ROI. Similarly, images of the same projection areas labelling MAP2, SV2A, and SYP were analyzed, and mean intensity values for the red channel were also measured in the defined ROIs.

Fiber colocalization of h-α-Syn with SERT and TH in projection areas was analyzed using the JACoP (Just Another Colocalization Plugin) plugin in ImageJ. First, images were standardized by adjusting their minimum and maximum pixel intensity values. Then, segmented images of the defined ROIs for the red and green channels were generated using consistent intensity thresholds across all images. Finally, the JACoP plugin was used to calculate Mander’s coefficient 1 (M1), which quantifies the proportion of the red channel overlapping with the green channel.

### Protein extraction and analysis by Western Blot (WB)

The procedure for WB and all antibodies used were previously validated in mouse models overexpressing human α-Syn and in postmortem human brain samples ^47,89,90^. Different brain areas from mice (mPFC, CPu, and HPC, 10 mg/each) and human (dlPFC, CPu, and HPC, 50 mg/each) were homogenized in RIPA lysis buffer (50 mM Tris, 150 mM NaCl, pH 8.0, 1% Triton X-100, 0.5% sodium deoxycholate, 5 mM EDTA, 0.1% SDS) with cOmplete™ protease and PhosSTOP™ phosphatase inhibitors (Roche; ref.: 05892970001, 4906845001, Basel, Sweden). Protein concentration was determined using the Pierce^TM^ BCA Protein Assay Kit (ThermoFisher Scientific, Waltham, MA, USA). 15 μg of protein lysates were separated by 4–15% SDS-PAGE (Bio-Rad, Hercules, CA, USA) and electrotransferred to a PVDF membrane (Bio-Rad). Protein blots were incubated with primary antibodies overnight at 4°C. Anti-h/m-α-Syn (1:1000; ref.: ab212184, Abcam), anti-SYP (1:1000; ref.: ab32127, Abcam), anti-SV2A (1:1000; ref.: PA5-110451, Fisher Scientific), anti-VAMP2 (1:2000; ref.: ab3347, Abcam), anti-SERT (1:1000; ref.: ab102048, Abcam), anti-PDS95 (1:3000; ref.: ab2723, Abcam), and anti-dynein (1:2000; ref.: MAB1618, Millipore) were used (**Table 2)**. Protein expression levels were normalized by using the housekeeping β-actin (anti-β-actin-HRP. 1:50000; A3854, Merck). ImageLab software (Bio-Rad) was used for analysis.

### Magnetic resonance imaging (MRI) acquisition, processing and analysis

Each mouse was scanned at 8 weeks after AAV2/5 vector (AAV-α-Syn or AAV-EV) infusion into DR (n=8 mice/group). Experiments were conducted on a 7.0T BioSpec 70/30 horizontal animal scanner (Bruker BioSpin), equipped with an actively shielded gradient system (600 millitesla per metre) with a surface coil for the mouse brain as receiver coil. Animals were sedated (4% isoflurane in 30% oxygen and 70% nitrogen), placed in a supine position in a Plexiglas holder with a nose cone for administering anesthetic gases and fixed using tooth and ear bars. Eyes were protected from dryness with Siccafluid ophthalmologic fluid. Once placed in the holder with constant isoflurane (1.5%), a subcutaneous bolus of medetomidine (Domtor, Orion Pharma; 0.3 mg/kg body weight) was injected. For the next 15 min, the percentage of isoflurane was progressively decreased to 0.5%. Then, a continuous perfusion of 0.6 mg/kg body weight per hour of medetomidine was started and maintained until the end of the acquisition session. After completion of the imaging session, 1 μl/g of atipamezole (Antisedan, Orion Pharma) and saline were injected to reverse the sedative effect and compensate fluid loss. Localizer scans were used to ensure the accurate position of the head at the isocentre of the magnet. Anatomical T2 RARE images were acquired in coronal orientation with effective time of echo (TE) = 33 ms, time of repetition (TR) = 2.3 s, RARE factor = 8, voxel size = 0.08 × 0.08 mm^2^ and slice thickness = 0.7 mm. Resting-state functional MRI (fMRI) was acquired with an EPI sequence with TR = 2 s, TE = 19.4, voxel size 0.21 × 0.21 mm^2^ and slice thickness = 0.5 mm. In total, 420 volumes were acquired resulting in an acquisition time of 14 min.

Seed-based analysis was performed as described previously ^91^ to evaluate functional connectivity of the CPu, HPC and mPFC with the rest of the brain. Preprocessing of the rs-fMRI data included: slice timing, spatial realignment using SPM8 for motion correction, elastic registration to the T2-weighted volume using ANTs for EPI distortion correction ^92^, detrend, smoothing (FWHM = 0.6 mm), frequency filtering of time series between 0.01 and 0.1 Hz, and regression by motion parameters using NiTime (http://nipy.org/nitime). Brain parcellation was performed based on the MRI-based atlas of the mouse brain ^93^. The atlas template was elastically registered to each subject T2-weighted volume using ANTs ^92^ and the estimated transformation was applied to the label map to obtain specific parcellation for each acquisition T2-weighted image, which was subsequently translated to the resting-state fMRI images. The left CPu, HPC or mPFC was selected as seed regions for seed-based analysis. Extracted average time series in the CPu, HPC or mPFC was correlated with the time series of all the voxels within the brain, resulting in a connectivity map for each seed, containing the value of the seed-to-voxel correlation. Voxel-wise statistics were computed in the seed-based connectivity maps using randomize algorithm from FSL ^94^ to identify areas where this connectivity was significantly different between groups. Familywise error (FEW) correction and TFCE (Threshold-free cluster enhancement) were applied to deal with multiple comparisons. Significance was defined as p < 0.05. The clusters of voxels where significant differences were identified were used as a mask to quantify average connectivity value in that area that was subsequently compared between groups. Together with this, for exploratory purposes, an additional voxel-based statistical analysis was performed including only the TFCE correction and setting a significance threshold of p < 0.005. If no significant differences were identified using FWE-correction p<0.05, results with uncorrected p < 0.005 will be reported. In that case, the clusters of voxels where significant differences were identified with this criterion will be also used as a mask to quantify average connectivity between the cluster and the seed and compared between groups.

### Statistical analysis

All values are mean ± standard error of the mean (SEM). Data from individual mouse was represented by single points when possible. The immunoreactivity value of the target proteins was corrected by the corresponding β-actin value and calculated as a percentage of a standard sample to control for inter-assay variability. The final estimate was the mean of the experimental replicates obtained in at least two different gels. Statistical comparisons were performed using GraphPad Prism 9.5.1 software. Outlier values were identified and excluded using the Grubbs’ test (i.e., Extreme Student zed Deviate – ESD method). Student’s *t* test, one-way and two-way ANOVA analysis, followed by Bonferroni *post hoc* test were performed when appropriate, and indicated in results and figure legends. Non-parametric Kruskal-Wallis test followed by Dunnett’s multiple comparison test was used to analyze differences in means between control and different stages of PD. Differences were considered significant if p < 0.05. Summary of the statistical analysis is shown in **Table 3**.

## ACKNOWLEDGEMENTS

This work was supported by MCIU/AEI/FEDER, UE grant (PID2019-105136RB-100, PID2022-141700OB-I00, MCIN/AEI/10.13039/501100011033 to A.B.), and AGAUR 2021-SGR-01358, Catalan Government. CB/07/09/0034 Center for Networked Biomedical Research on Mental Health (CIBERSAM). We also thank the Spanish Stress Research Network, MCIN/AEI /10.13039/501100011033. U.S.S. is a recipient of a fellowship from the Non-Doctor Researcher Formation Pre-doctoral Program of Basque Government, Spain. U.A. is a recipient of a fellowship from the State Research Agency for the training of predoctoral research personnel (PREP2022-000676) associated to a generation of knowledge project (PID2022-141700OB-I00). The authors would like to thank the staff members of the Biobank of the Clinic Hospital Barcelona - IDIBAPS for their collaboration in the study.

## AUTHOR CONTRIBUTIONS

Conceptualization and supervision of the research: A.B. Methodology: L.M.R. and U.S.S. performed stereotaxic surgeries in mice; L.M.R. processed human brain tissue samples; L.M.R., J.J.E, U.S.S., C.Y.C., U.A., and M.T.L. performed immunohistochemistry and WB experiments; V.P. performed *in situ* hybridization; C.C. provided advice on the acquisition and analysis of images on the Dragonfly confocal microscopy system; E.M.M. and X.L.G. conducted the rs-fMRI experiments, image acquisition, and data analysis. L.M.R. analyzed and supervised the data. Writing: A.B. with all nine authors’ inputs. Funding acquisition: A.B. All authors edited and approved the final version of the manuscript.

## ADDITIONAL INFORMATION

Supplementary information

## Figure legends

**Figure Suppl 1. Overexpressing of h-α-Syn in raphe 5-HT neurons of female mice.** Female mice received 1 μl of AAV2/5-CBh-WPRE3 construct to drive expression of h-α-Syn (AAV-h-α-Syn) or empty AAV2/5-CBh-WPRE3 vector containing non-coding (null) stuffer DNA (AAV-EV) into dorsal raphe nucleus (DR) and were euthanized at 4 and 8 weeks (W) post-injection. **A**) Representative confocal image showing co-localization of h-α-Syn protein and TPH^+^ cells in the DR of mice at different time points after AAV2/5 injection. Scale bar: low: 200 μm. On the right, schematic representation of h-α-Syn^+^/TPH^+^ cell density. **B)** Bar chart (top) showing progressive accumulation of h-α-Syn protein in mouse DR. Bar chart (down) showing the proportion of the total number of TPH^+^ cells co-localized with h-α-Syn^+^ (n = 5 mice/group; ***p< 0.0001 versus AAV-EV mice). Values are presented as mean ± SEM.

**Figure Suppl 2. Widespread h-α-Syn transport to 5-HT projection brain areas.** Male mice received 1 μl of AAV2/5-CBh-WPRE3 construct to drive expression of h-α-Syn (AAV-h-α-Syn) or empty AAV2/5-CBh-WPRE3 vector containing non-coding (null) stuffer DNA (AAV-EV) into dorsal raphe nucleus (DR) and were euthanized at 4 and 8 weeks (W) post-injection. **A**) Representative confocal image showing co-localization of h-α-Syn^+^/SERT^+^ fibers in 5-HT projection brain regions, such as medial prefrontal cortex (mPFC), cingulate cortex (Cg), caudate putamen (CPu), and hippocampus (HPC) of mice at different time points after AAV2/5 injection. Scale bar: 25 μm. **B)** Bar charts show the density of SERT^+^ fibers in the brain areas analyzed: prelimbic cortex (PrL), infralimbic cortex (IL), Cg, CPu, and different HPC subfields (n = 5 mice/group; * p< 0.05, **p< 0.01, ***p< 0.001 versus AAV-EV or AAV-h-α-Syn 4W mice). Values are presented as mean ± SEM.

**Figure Suppl 3. Axonal transport of h-α-Syn in TH-positive fibers originating in the DR.** Male mice received 1 μl of AAV2/5-CBh-WPRE3 construct to drive expression of h-α-Syn (AAV-h-α-Syn) or empty AAV2/5-CBh-WPRE3 vector containing non-coding (null) stuffer DNA (AAV-EV) into dorsal raphe nucleus (DR) and were euthanized at 4 and 8 weeks (W) post-injection. **A, B**) Representative confocal image showing co-localization of h-α-Syn^+^/TH^+^ fibers in cingulate cortex (Cg) and caudate putamen (CPu) of mice at different time points after AAV2/5 injection. Scale bar: 25 μm. **C, D)** Bar chart (left) showing progressive accumulation of h-α-Syn protein in mouse Cg and CPu, respectively. Bar chart (right) the density of TH^+^ fibers in the brain areas analyzed (n = 5 mice/group; ***p< 0.01 versus AAV-EV mice). Values are presented as mean ± SEM.

