## Supplemental Figures for "Early synaptic changes and reduced brain connectivity in PD-like mice with depressive phenotype"

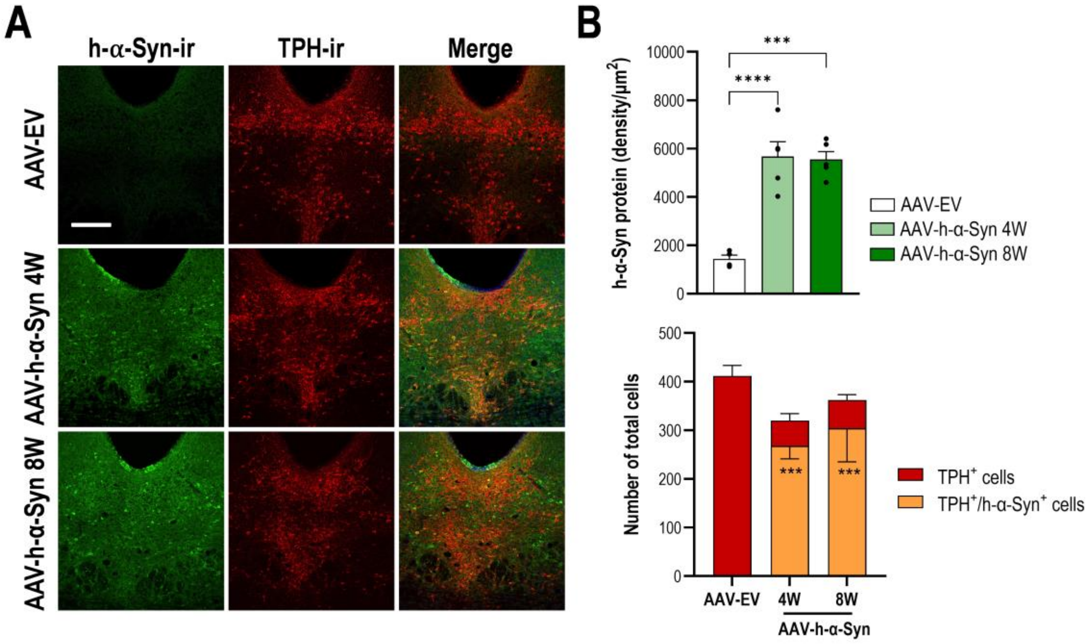


**Figure Suppl 1.**

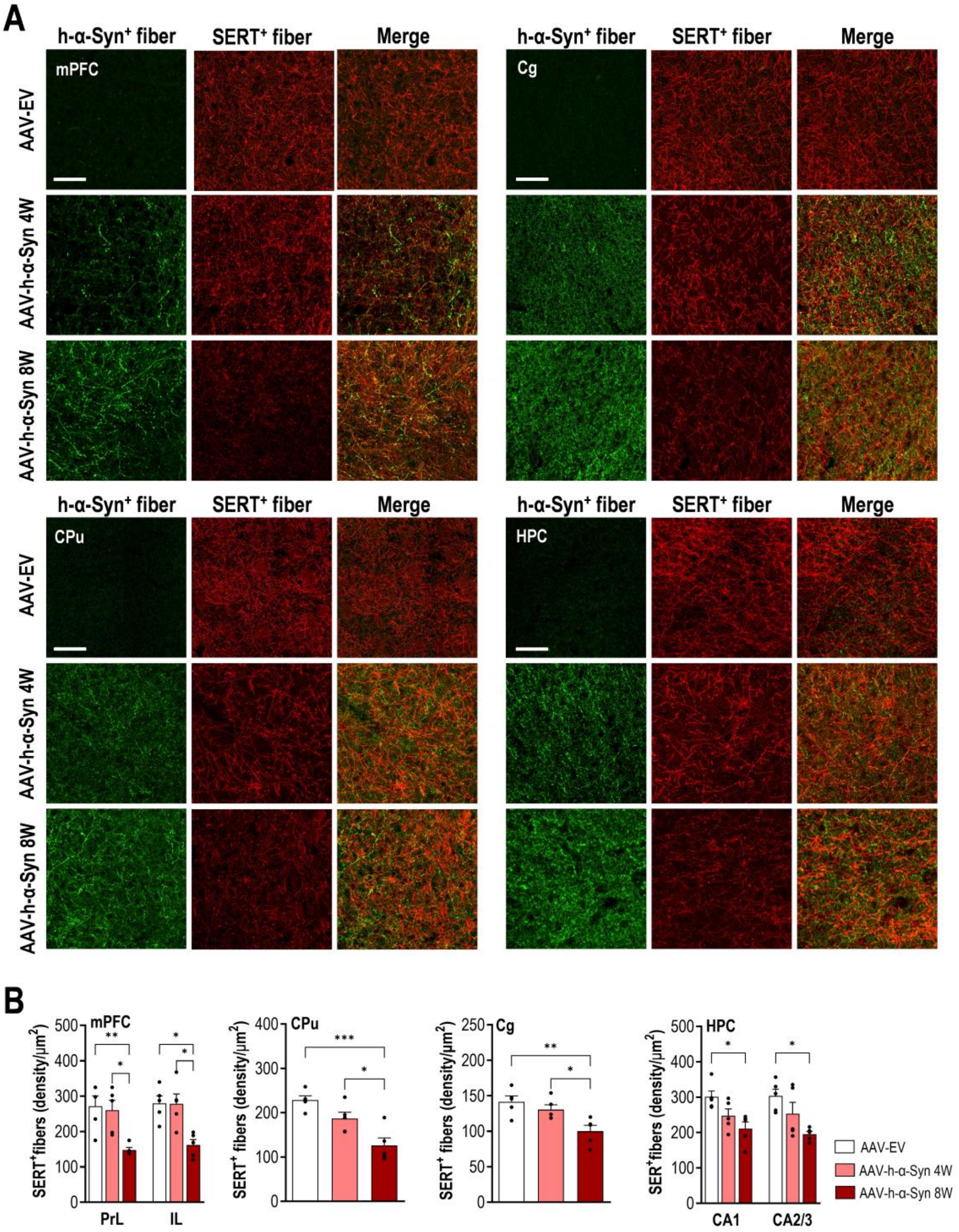


**Figure Suppl 2.**

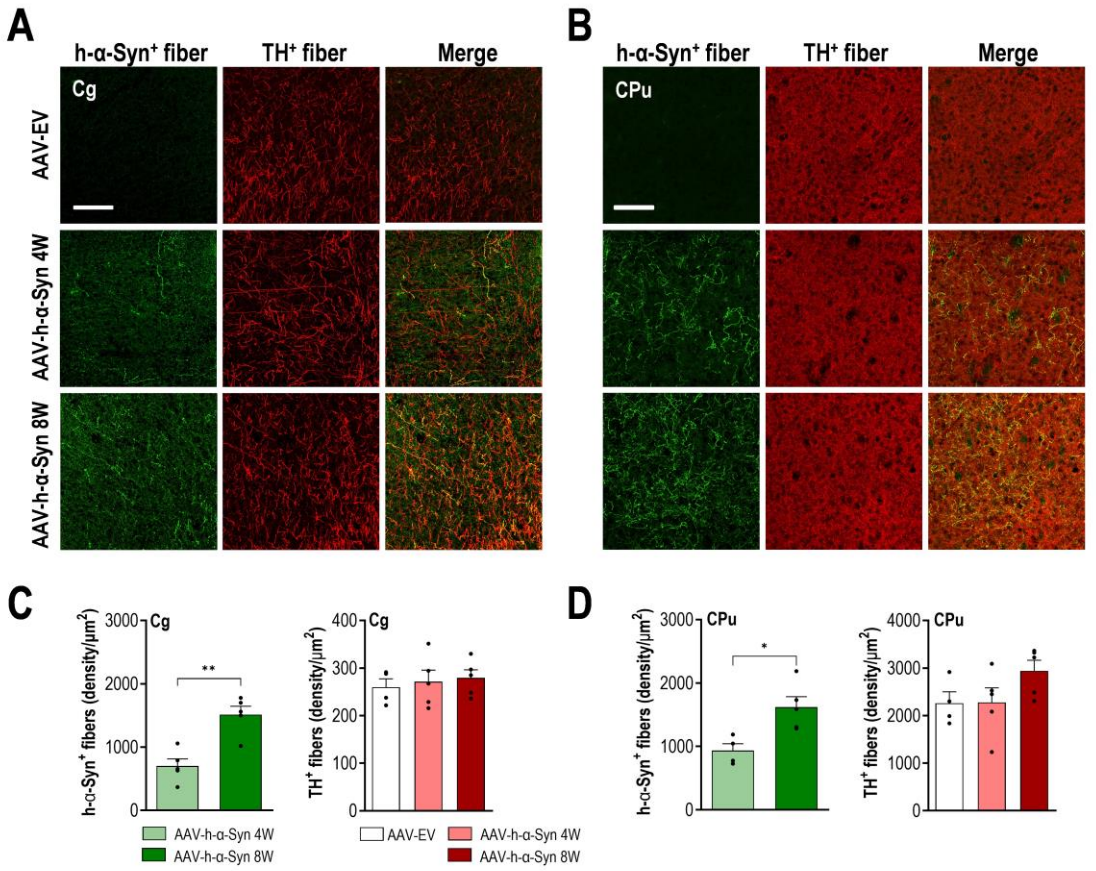


**Figure Suppl 3.**
